# scRegulate: Single-Cell Regulatory-Embedded Variational Inference of Transcription Factor Activity from Gene Expression

**DOI:** 10.1101/2025.04.17.649372

**Authors:** Mehrdad Zandigohar, Jalees Rehman, Yang Dai

## Abstract

**Motivation:** Accurately inferring transcription factor (TF) activity from single-cell RNA sequencing (scRNA-seq) data remains a fundamental challenge in computational biology. While existing methods rely on statistical models, motif enrichment, or prior-based inference, they often depend on deterministic assumptions about regulatory relationships and rely on static regulatory databases. Few approaches effectively integrate prior biological knowledge with data-driven inference to capture novel, dynamic, and context-specific regulatory interactions.

**Results:** To address these limitations, we develop scRegulate, a generative deep learning framework leveraging variational inference to estimate TF activities guided by experimental TF-target gene relationships and progressively adapted based on the input scRNA-seq data. By integrating structured biological constraints with a probabilistic latent space model, scRegulate offers a scalable and biologically grounded estimation of TF activity and gene regulatory network (GRN). Comprehensively bench-marking on public experimental and synthetic datasets demonstrates scRegulate’s superior ability. Further, scRegulate accurately recapitulates experimentally validated TF knockdown effects on a Perturb-seq dataset for key TFs. Applied to experimental human PBMC scRNA-seq data, scRegulate infers cell-type-specific GRNs and identifies differentially active TFs aligned with known regulatory pathways. scRegulate’s TF activity representations capture transcriptional heterogeneity, enabling accurate clustering of cell types. scRegulate is highly efficient, frequently an order of magnitude faster than common baselines. Collectively, our results establish scRegulate as a powerful, interpretable, and scalable framework for inferring TF activities and GRNs from single-cell transcriptomics.

**Availability:** Results and scripts available at github.com/YDaiLab/scRegulate.

**Supplementary information:** Supplementary data are available at *Bioinformatics* online.

## 1 Introduction

Transcription factors (TFs) drive cellular identity and function by regulating gene expression (Lambert *et al*., 2018; Voss and Hager, 2014). When TFs bind to specific DNA motifs and recruit regulatory machinery, they modulate the activation or repression of target genes in a context-dependent manner. Precise regulation of their activity is essential not only for normal cellular processes but also for survival, development, and response to environmental cues. Conversely, their dysregulation can result in a wide range of diseases. Accurate inference of TF activity is thus essential for understanding cell fate decisions and cell state transitions during key biological processes, as well as disease development and progression.

Single-cell RNA sequencing (scRNA-seq) has revolutionized the study of gene regulation by enabling high-resolution profiling of transcriptomes in individual cells. Numerous computational methods exist for inferring TF activity from single-cell transcriptomes (Hecker *et al*., 2023). decoupleR (Badia-i-Mompel *et al*., 2022) rapidly estimates TF activity by coupling a prior knowledge network of TF-target genes with statistical methods, such as the univariate linear model, or an ensemble of different models. While it offers superior speed advantages due to its simplicity, its capacity is limited in capturing complex regulatory relationships. SCENIC integrates known TF-target gene databases and motif enrichment analysis to infer the activity of a TF regulon (i.e., target genes of the TF) based on co-expression that may not be necessarily expected between the TF and a target gene (Aibar *et al*., 2017). In addition, its dependence on static motif databases for pruning may limit its accuracy when inferring TF regulon activity in poorly annotated systems. Bayesian factor models, such as BITFAM (Gao *et al*., 2021), bypass co-expression and motif-based inference by incorporating prior knowledge of TF binding data retrieved from public TF-ChIP databases. The probabilistic matrix factorization model in BITFAM results in two key outputs: (1) inferred TF activities in individual cells, and (2) a weighted TF-target gene matrix. This matrix constitutes a weighted gene regulatory network (GRN), a definition that will be used in this work. However, the reliance on linear assumptions may still limit their ability to capture the complex non-linear transcriptional dynamics present in heterogeneous cell populations.

Multiomic approaches, such as SCENIC+ (Bravo González-Blas *et al*., 2023), CellOracle (Kamimoto *et al*., 2023), and BIOTIC (Cao *et al*., 2025) attempt to enhance inference by integrating chromatin accessibility alongside single cell transcriptomic data. These methods can provide more accurate predictions of TF-target gene relations. However, they require high-quality enhancer annotations or complementary ATAC-seq chromatin data, which are not always available for scRNA-seq datasets generated under specific biological conditions. Additionally, their reliance on integrating diverse modalities introduces additional complexity and presents significant challenges data harmonization.

GRNs and TF activities represent crucial regulatory mechanisms in dnamic biological processes and disease progression. GRNs are inherently context-dependent and cell-type-specific, necessitating models for GRN inference that can adapt dynamically to transcriptional profiles of different cell types or cell states. However, the inherent sparsity and noise in scRNA-seq data may adversely impact the accuracy of the inference of TF activity and GRN. In light of these challenges, recent developments in generative models, particularly variational autoencoders (VAEs), have shown promise in extracting meaningful representations from high-dimensional scRNA-seq data (Lopez *et al*., 2020; Gayoso *et al*., 2022). While models like SCALE (Xiong *et al*., 2019) and other adaptations of scVI (Svensson *et al*., 2020) have made the decoder structure more interpretable, they are not designed for inference of TF activity and GRN.

To address these challenges, we develop scRegulate, a novel framework that explicitly embeds a prior GRN in a VAE architecture. Specifically, scRegulate integrates curated TF-target interactions from existing regulatory datasets into a non-linear generative model at the start of inference. Unlike previous approaches that infer static or bulk-level GRNs, scRegulate dynamically tailors TF-target relationships and TF activities (i.e., the activation of TF nodes in the VAE) to reflect the unique transcriptional landscapes of different cell populations. This capability provides unprecedented resolution in studying cell-type-specific regulatory mechanisms and their roles in cellular identity and function, allowing for dynamic refinement of TF-target interactions. Through comprehensive benchmarking, we demonstrate that scRegulate outperforms existing methods in accuracy, robustness, and biological relevance.

## 2 Materials and Methods

### 2.1. Overview

scRegulate is a GRN inference framework that integrates prior knowledge of TF and target genes into a VAE architecture (Figure 1). The model follows a structured three-phase approach: (1) in the prior initialization phase, the weights connecting the TF activity layer to the output layer are initialized to enforce known TF-gene regulatory interactions; (2) in the dynamic inference phase, regulatory weights are optimized, with prior constraints gradually relaxed to allow for the discovery of new TF-target interactions; and (3) in the cell-type-specific fine-tuning phase, GRNs and TF activities are optimized per cell-type (or per cell cluster) for more accurate representation of cell-type-specific regulation. In scRegulate, TF activities are defined by the cell-specific TF representations learned from the VAE (Figure 1), and are computed from the ReLU activations. Thus, TF activities are non-negative values, where higher values indicate stronger relative regulatory effects of a TF on a given cell. In contrast, an inferred GRN weight matrix contains positive, negative, or zero values. Here, the sign indicates whether the TF functions as an activator (positive) or repressor (negative) of a target gene, while the magnitude reflects the relative strength of regulation. To facilitate interpretation, we provide both the raw GRN weights and post-processed versions obtained by min–max scaling each TF’s outgoing weights to the range [0,1]. The scaled weights enable relative comparison across TFs and conditions and were used in our benchmarking analyses. Together, TF activities provide a cell-specific readout of regulatory influence, while GRN weights describe the direction and strength of TF-gene interactions. The inferred TF-activities and weighted GRNs can be used for downstream analysis, including identifying differentially active TFs, cell-type-specific TF-gene interactions, and functional enrichment analyses for the target genes. The three core modules in the scRegulate pipeline, i.e., data preprocessing, TF activity inference via VAE, and downstream analysis, are implemented using Scanpy (Wolf *et al*., 2018), adhering to best practices in scalable single-cell omics analysis.

**Figure 1.**
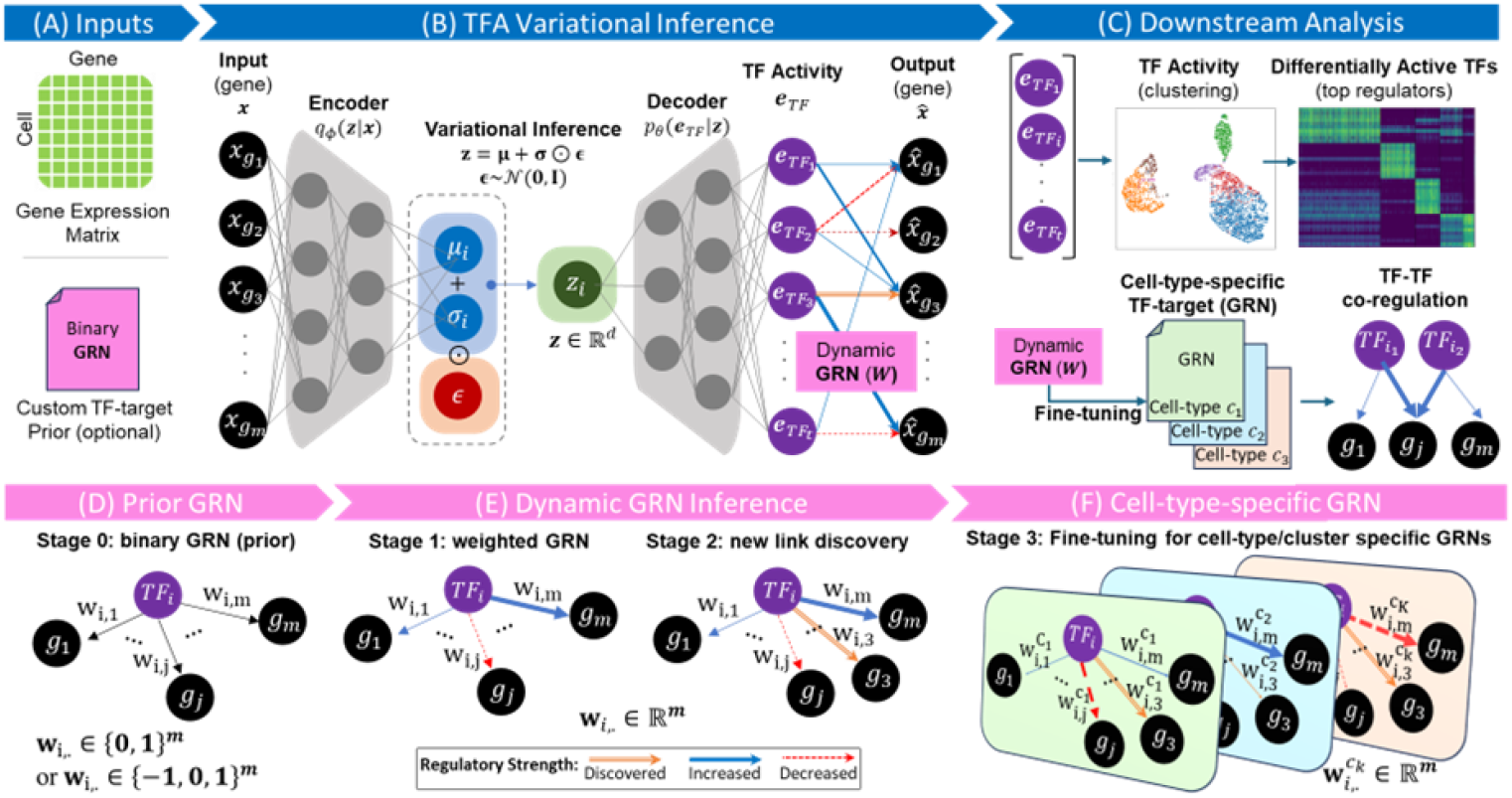
Overview of scRegulate for TF activity and GRN inference. **(A)** Inputs include a gene expression matrix and a TF-target gene prior GRN. **(B)** TF activity is inferred through a variational autoencoder framework, where the encoder maps input data into a latent space (**z**), followed by a TF activity representation layer (***e***_*TF*_) to predict gene expression and capture TF activities, informed by the dynamic GRN. **(C)** Downstream analysis leverages TF activities (***e***_*TF*_) for clustering, identification of differentially active TFs. Construction of cell-type-specific weighted GRNs (***W***), provides insights into TF-target regulation and TF-TF co-regulation. **(D)** Prior GRN initializes the regulatory network using a binary or ternary TF-target prior. **(E)** Dynamic GRN inference refines the network by adjusting regulatory strengths (edge weights of TF-target gene links), discovering new connections, and evolving to a weighted GRN. **(F)** Fine-tuning the GRN for individual cell types or clusters, resulting in cell-type-specific regulatory networks. Notations are summarized in Table S1.

### 2.2. Data Preprocessing

The input data consists of a scRNA-seq expression matrix and an optional prior GRN. The CollecTRI (Müller-Dott *et al*., 2023), a high-quality collection of signed TF-gene interactions for 1186 TFs, is leveraged as default to initialize our regulatory network. The scRNA-seq data undergoes standard preprocessing using Scanpy, including filtering out low-quality cells based on mitochondrial gene content, selecting highly variable genes, and performing log-normalization. Further filtering is applied such that only genes expressed in at least 10 cells and cells expressing at least 3 genes are retained. Additionally, genes must have at least one associated TF in the prior GRN to be included. TFs regulating fewer than 10 target genes in the data are removed from consideration. These preprocessing steps are intended to reduce the risk of inferring spurious regulatory links from TFs with limited target information.

### 2.3. Model Architecture and Implementation

The VAE architecture consists of fully connected layers with ReLU activation functions, forming both encoder and decoder networks. The encoder (1-3 layers) maps input gene expression profiles to a lower-dimensional latent space, while the decoder (1-2 layers) first infers TF activity and then reconstructs gene expression using a trainable GRN. Latent variables are inferred via the reparameterization trick, and the model is trained using variational inference. Optimization details, including loss functions and hyperparameter selection, are described in Section 2.3.4. The VAE model was implemented using PyTorch (Paszke *et al*., 2019).

#### 2.3.1. Variational Inference of TF Activity

scRegulate employs a VAE to infer latent TF activity from gene expression data. The input gene expression vector **x** from a cell is mapped to a latent representation ***z*** using an encoder network parameterized by mean **μ** and variance **σ**:

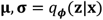

here, *q*_*ϕ*_(**z** ∣ **x**) denotes the encoder network, which outputs the mean ***μ*** and standard deviation **σ** of the latent variable **z** given the input **x**.

The latent variable **z** is obtained by shifting and scaling a standard normal random variable **ϵ** drawn from *N*(**0, I**), also known as the reparameterization trick:

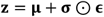

The latent representation **z** is decoded by *p*_*θ*_ to estimate TF activities ê_TF_:

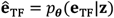

here, **e**_TF_ represents the unobserved true TF activities that the model aims to estimate. These inferred TF activities are then mapped to gene expression through the GRN weight matrix **W**:

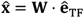

where 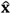 is the reconstructed gene expression vector, ê_TF_ is the inferred TF activity vector (or learned TF representations), and **W** is the weight matrix defining TF-gene interactions, i.e., the weighted GRN. This explicit relationship defined by the last layer in the neural network is one of the key features of scRegulate, allowing direct examination of TF effects on target gene expression and enhancing model interpretability.

#### 2.3.2. Integration of Prior GRNs into TF Activity Estimation

To initialize ê_TF_, scRegulate employs a Univariate Linear Model (ULM) inspired by (Badia-i-Mompel *et al*., 2022), where TF activities are estimated as a weighted sum of all target gene expression values in the prior GRN (Supplementary Note 1). To balance the contribution between the prior-driven estimate 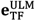 and the data-driven estimate *p*_*θ*_(**e**_TF_ |**z**) from decoder, the contribution of 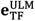 is gradually reduced over training by a dynamic scheduling parameter *α*:

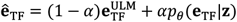

where *α* ∈ [0, *α*_*max*_], and *α* follows an increasing schedule to transition from prior-driven 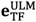 toward data-driven inference *p*_*θ*_(**e**_TF_|**z**). Further, the scheduling of *α* follows a controlled incremental approach, transitioning from its initial value *α*_start_ to its maximum *α*_max_ in discrete steps defined by the hyperparameter Δ*α*. The update rule for *α* is formulated as:

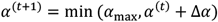

here, each training iteration is denoted as *t*.

#### 2.3.3. GRN Prior Enforcement and Dynamic Updating

To reduce noise in the inferred regulatory interactions, weak regulatory interactions in the **W** are penalized by enforcing sparsity as L1 regularization:

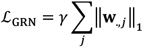

where *γ* is the regularization parameter linearly scheduled over training epochs, and **w**_.,**j**_ is the vector of regulatory weights for gene **j**. The prior GRN, **W**_prior_, is enforced using a mask factor *m*(*t*), which gradually transitions from 1 to 0 over training epochs:

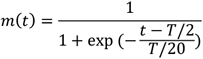

where *T* is the total number of epochs. This ensures that initial regulatory constraints are respected while allowing for new regulatory interactions to emerge. The mask factor is applied to regulate updates to:

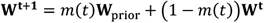

This updates the GRN weights by blending the previous weight matrix with the newly learned weights, modulated by the mask *m*(*t*), ensuring a smooth transition from prior-based regulation to learned interactions.

#### 2.3.4. Loss Functions, Optimization, and Training Details

The VAE is trained to minimize the total Loss which is the sum of the Evidence Lower Bound (ELBO) and the GRN regularization loss:

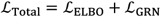

To ensure model generalizability and prevent overfitting to noise, we apply an 85%-15% split between the training and validation subsets during model training. The optimizer used for training is Adam, with an adaptive learning rate scheduler that dynamically adjusts based on validation loss. More details regarding the training are available in the Supplementary Note 2.

For benchmarking runs, fine-tuning was performed globally across all cells in an unsupervised manner, without using any cell-type labels for fairness.

#### 2.3.5. Benchmark Datasets

Multiple datasets were used to evaluate clustering of inferred TF activities, GRNs, and biological validity of TF activities. To investigate how well the inferred TF activities capture the cell heterogeneity in the data, we used a combination of publicly available scRNA-seq datasets where ground-truth cell-type labels are available. We used three Tabula Muris datasets (heart, lung, and brain) (The Tabula Muris Consortium *et al*., 2018), PBMC, and human middle temporal gyrus (MTG) scRNA-seq (Allen Institute for Brain Science, 2022) (Supplementary Table S2).

For GRN benchmarking, we utilized the GRouNdGAN (Zinati *et al*., 2024) simulated single-cell datasets with imposed regulatory interactions, making it a controlled benchmark for testing network reconstruction accuracy (Supplementary Note 3 & Table S3). To ensure robust performance evaluation and stability across different subsets, we employed proportional stratified sketching to generate five independent subsets. This approach ensures that each subset retains the full diversity of cell types while maintaining proportional representation across the dataset, reducing bias introduced by random sampling. The procedure involved clustering the Leiden algorithm for clustering, followed by sketch-based subsampling within each cluster using geosketch. Each pseudo-cluster was assigned a subset size proportional to its overall representation in the dataset. This method ensures that smaller clusters were not underrepresented while maintaining a well-balanced distribution across all five subsets. Additionally, splitting the dataset into multiple subsets provides a bounded range for GRN performance metrics, ensuring that results are not biased by a single instance of dataset partitioning.

For TF activity benchmarking, we employed a Perturb-seq dataset, in which specific TFs were inhibited via CRISPR interference (CRISPRi) (Dixit *et al*., 2016). The experimental TF knockdown effects were used as a reference for evaluating inferred TF activities.

Finally, to demonstrate real-world applicability, we applied scRegulate to a PBMC 3k dataset (Supplementary Table S2), a well-characterized immune cell dataset. This dataset comprises peripheral blood mononuclear cells from healthy donors. This allowed for an in-depth exploration of GRN structures and TF activity inference at single-cell resolution, with extensive analysis of TF-TF interactions, differentially active TFs, and functional gene enrichment analysis.

### 2.4. Benchmarking Strategy and Evaluation Metrics

Benchmarking was performed against five representative baseline tools (pySCENIC, BITFAM, decoupleR, BIOTIC, CellOracle) chosen as top-performing and widely used methods. Supplementary Table S4 lists additional GRN inference tools that were not included in our comparisons but could be of value in certain contexts. For benchmarking, BITFAM was run with its default GTRD prior. SCENIC, CellOracle, and BIOTIC were used following their recommended pipelines. scRegulate and decoupleR were initialized with CollecTRI as default.

#### 2.4.1. Clustering Performance Evaluation

Leiden clustering is applied to the inferred TF activities at multiple resolutions (0.2-3.0), and cluster assignments are compared to ground truth labels using standard clustering metrics. Each cluster is assigned a label based on maximum voting. Specifically, for each cluster, we tally the ground-truth cell-type labels of all the cells within that cluster. The label that appears most frequently is then assigned as the representative label for the cluster. This process enables the computation of clustering evaluation metrics (ARI, NMI, macro F1) against the known cell-type annotations. Formal definitions and computational formulas for these evaluation metrics are provided in Supplementary Note 4. To assess the robustness of clustering using inferred TF activities, we simulated dropout effects by applying a masking process where each gene expression value *X*_*ij*_ was retained with probability 1 − *p* and set to zero otherwise, using:

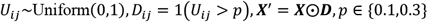

This mimics the sparsity commonly observed in scRNA-seq data.

#### 2.4.2. GRN Inference Evaluation

GRN inference performance is evaluated using AUROC and AUPRC, computed over the intersection of target genes between scRegulate-inferred networks and the ground-truth networks in GRouNdGAN. The inferred weighted GRNs are binarized at varying thresholds and compared against a ground-truth binary GRN to assess performance stability. The GRN consistency across cell types was evaluated by the Pearson correlation between TF-target matrices after applying min-max normalization based on absolute values. Hierarchical clustering on the resulting correlations of cell-type-specific GRNs was performed to identify groups of cell types with similar GRNs (Supplementary Note 5).

#### 2.4.3. Benchmarking TF Activity using the Real Perturb-seq Data

The performance of scRegulate in inferring TF activity was evaluated using a Perturb-seq dataset, where experimental perturbations (i.e., knockdowns) of key TFs were performed. Predicted TF activities were compared between perturbed and control groups by averaging within each condition and computing the *log*_2_ fold change (LFC) between conditions. In addition, the Wilcoxon rank-sum tests were performed to assess the capability of scRegulate to distinguish perturbed TFs from non-perturbed ones.

### 2.5. Application of scRegulate to the Experimental Human PBMC Data

We illustrated the utility of scRegulate on the experimental human PBMC 3k dataset, which was first preprocessed following the standard scRNA-seq pipeline by Scanpy. scRegulate was then used to infer cell-type-specific TF activities and reconstruct GRNs at single-cell resolution. The results were then investigated for comparative and functional biological interpretations.

#### 2.5.1. Comparative Analysis of the Inferred TF Activity

We performed a differential analysis of TF activities across cell types. Specifically, we applied Wilcoxon rank-sum test to compare TF activity profiles between groups, retaining only those TFs with an adjusted p-value below 0.05. From this subset, we ranked the differentially active TFs based on the highest LFC to prioritize the most robustly modulated regulators.

#### 2.5.2. Functional Validation of the GRN

We focused on validating the cell-type-specific GRNs inferred by scRegulate within the PBMC dataset. For this case study, we applied cell-type-specific fine-tuning procedure, where GRNs were optimized separately within each annotated cell type (Note: the find-tunning does not require annotation and can be applied to unannotated cell clusters). We evaluated the similarity of GRNs across cell types by computing Pearson correlation coefficients (described in Section 2.4.2) to identify groups of cell types with similar GRNs. To validate the inferred GRNs, we performed Gene Ontology (GO) enrichment analysis by ranking each TF’s target genes based on GRN weights. Enrichment was evaluated using the Azimuth Cell Types 2021 gene sets, applying an adjusted p-value threshold of 0.05 and selecting the top 1% of targets. Relevant GO terms were curated for downstream analyses. In parallel, a TF co-regulatory network was constructed by linking TFs with high cosine similarity in their regulatory profiles, highlighting modules of coordinated transcriptional regulation across cell types. A detailed description of the TF-TF co-regulation similarity analysis, including cosine similarity calculations across GRN weight vectors, is provided in Supplementary Note 6.

## 3 Results

### 3.1. The Inferred TF Activities Capture Cellular Heterogeneity

scRegulate’s inferred TF activities provide a structured representation of cellular heterogeneity. Clustering analyses revealed that the TF activities inferred from scRegulate achieved highly accurate cell-type separations, surpassing other TF-based methods in key clustering metrics (Figures 2A, and S1). For PBMC, scRegulate achieved an ARI of 0.814 ± 0.042, NMI of 0.821 ± 0.021, F1-score of 0.868 ± 0.118, and AUC of 0.930 ± 0.058, significantly outperforming SCENIC, decoupleR and BIOTIC (Wilcoxon signed-rank test, p < 0.01). Similar trends were observed across the other datasets, with scRegulate achieving an AUC of 0.948 ± 0.033 in lung, 0.992 ± 0.002 in heart, and 0.953 ± 0.015 in brain, outperforming or competing with the other methods.

**Figure 2.**
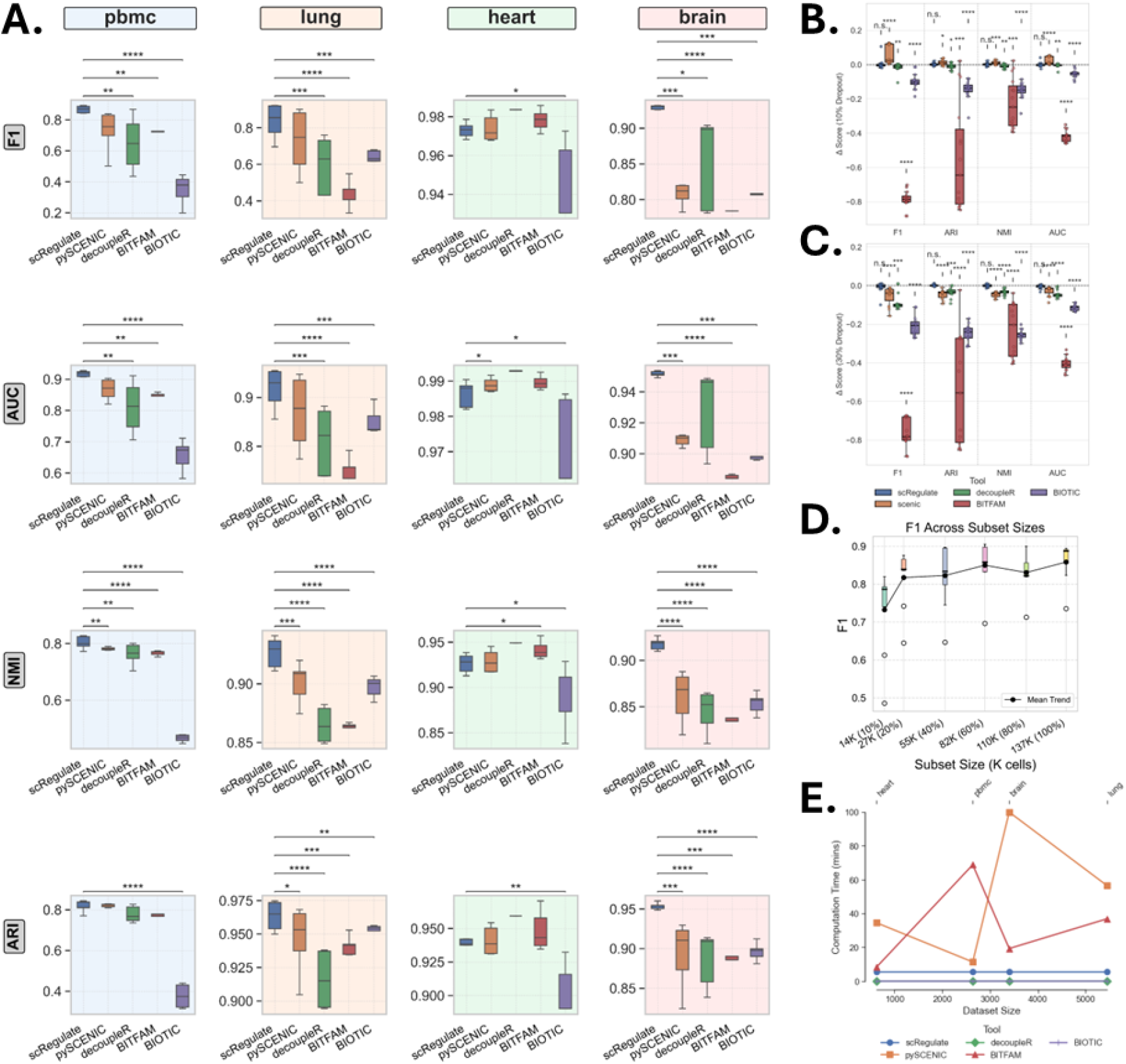
Comparison of clustering using TF activity-based inferred from the five different methods on four scRNA-seq datasets. **(A)** Clustering performance (ARI, NMI, F1, and AUC) on using TF activities inferred from four methods (scRegulate, SCENIC, BITFAM, decoupleR and BIOTIC) using the PBMC, mouse brain, mouse lung, and mouse heart datasets. Each boxplot displays performance using different Leiden clustering resolutions; asterisks indicate statistical significance based on p-values (p < 0.05: *; p < 0.01: **; p < 0.001: ***). **(B)** Effect of 10% simulated dropout and **(C)** 30% simulated dropout on using the brain Tabula Muris dataset, highlighting each method’s robustness to partial TF information. **(D)** F1-score across dataset sizes for the MTG dataset, showing performance scalability. **(E)** Computation times for each method across the four datasets, illustrating differences in runtime as dataset size and complexity vary. In all plots, error bars or box boundaries represent variability across different Leiden resolutions.

To assess the robustness of clustering based on scRegulate’s inferred TF activity, we simulated dropout effects by randomly removing 10% and 30% of gene expression values in the MTG dataset, mimicking the sparsity commonly observed in single-cell data (Figures 2B, C and S2). scRegulate maintained high clustering performance across different Leiden resolutions, whereas the alternative methods exhibited significant performance degradation (Wilcoxon signed-rank test, p < 0.01). scRegulate achieved consistently high F1-scores as dataset size increased, demonstrating its scalability for large-scale scRNA-seq datasets (Figures 2D, S3). Finally, we evaluated computation times across all methods and observed that scRegulate efficiently scales with dataset size, outperforming SCENIC and BITFAM in runtime while maintaining high accuracy (Figure 2E). Taking together, these results highlight scRegulate’s ability to generate biologically meaningful TF activity representations that more accurately reflect transcriptional variation across cell types, reinforcing its utility for robust single-cell clustering.

### 3.2. scRegulate Outperforms Existing Methods in GRN Inference on Synthetic and Experimental Gold Standard data

Across all benchmarked synthetic datasets (Dahlin, PBMC, and Tumor), scRegulate consistently outperformed SCENIC, CellOracle and BITFAM in GRN inference based on AUROC and AUPRC metrics (Figures 3A, and S4). Notably, in the PBMC dataset, scRegulate achieved the highest AUPRC (0.946) and a strong AUROC (0.703), significantly surpassing pySCENIC (AUPRC = 0.746, AUROC = 0.568), CellOracle (AUPRC = 0.375, AUROC = 0.549), and BITFAM (AUPRC = 0.398, AUROC = 0.537). A similar trend was observed in the Tumor dataset, where scRegulate’s AUPRC reached 0.840, exceeding pySCENIC (0.744), CellOracle (0.516), and BITFAM (0.490). The Dahlin dataset followed this pattern, with scRegulate achieving an AUPRC of 0.796, while pySCENIC, CellOracle, and BITFAM scored lower at 0.588, 0.286 and 0.313, respectively. From a mouse embryonic stem cell (mESC) scRNA-seq dataset (Table S2), we benchmarked scRegulate against pySCENIC and observed improved AUROC and AUPRC based on the evidence from ChIP, Perturb, and ChIP+Perturb (McCalla *et al*., 2023) with markedly shorter runtimes (Fig. S5).

**Figure 3.**
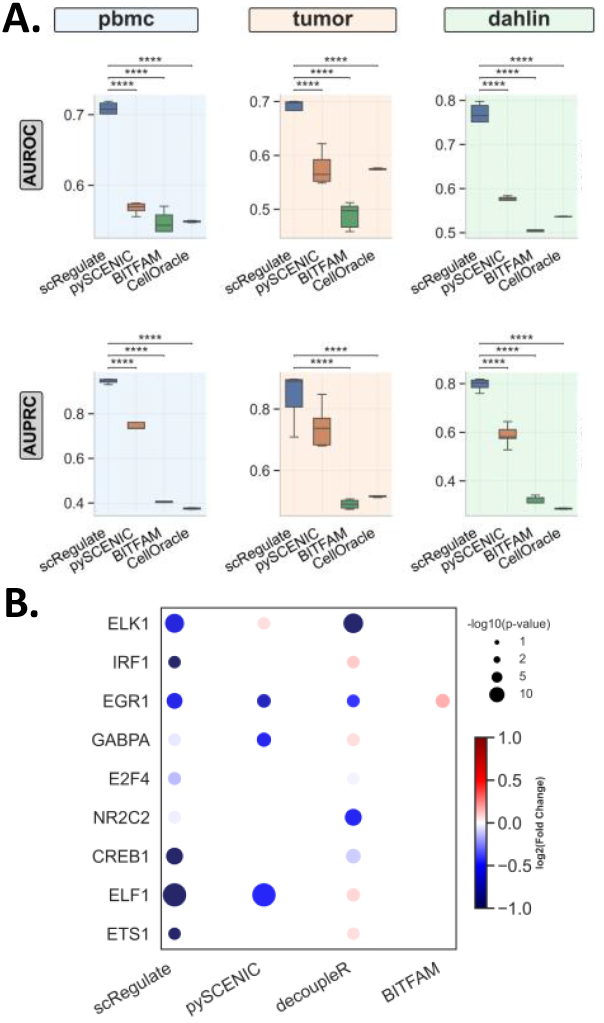
Comparison of scRegulate with Alternative Tools on GRN Inference. **(A)** GRNs inferred from the three synthetic ground-truth data of PBMC, tumor, and Dahlin. **(B)** TF activity inferred using real perturb-seq dataset with double knockdown effect (****: p < 0.0001).

### 3.3. scRegulate Outperforms Alternatives in Predicting TF Activity from Experimental Knockdowns

To assess scRegulate’s ability to correctly infer TF activity, we evaluated its performance on the Perturb-seq dataset. Since CRISPRi leads to transcriptional repression, we expect negative log-fold changes (LFCs) in the inferred TF activities when comparing the knockdown cells vs. control cells. We focused on double-knockdown perturbations to minimize off-target effects and provide a more reliable benchmarking framework for TF activity inference. The dataset comprised double-knockdown perturbations for 10 TFs, with all available TFs included by selecting cells expressing multiple gRNAs per TF to ensure robust functional knockdown (Table S2). Across all tested perturbations, scRegulate consistently inferred significantly reduced TF activity in the knockdown cells compared to controls (Wilcoxon signed-rank test, p < 0.01). TF activity log-fold changes among significantly repressed TFs ranged from –0.19 to –0.61, with ELK1 (LFC = –0.61, p = 2.37×10−^11^) and EGR1 (LFC = –0.57, p = 7.12×10−^6^) showing the strongest repression. Moderate decreases were also observed for TFs such as CREB1 (LFC = –0.28, p = 0.011), confirming that scRegulate correctly captured expected repression patterns (Figures 3B and S6). Compared to the alternative methods, scRegulate demonstrated higher precision in detecting TF knockdowns. For ELK1, both SCENIC and decoupleR failed to detect significant repression (p > 0.7), while BITFAM incorrectly inferred increased activity (LFC = 0.18, p = 0.0068). In contrast, scRegulate’s results were more consistent with ground truth expectations. These results confirm that scRegulate provides an accurate and biologically consistent framework for inferring TF activity from the CRISPRi-based Perturb-seq data, correctly capturing repression effects where other methods struggled.

### 3.4. scRegulate Infers Cell-Type-Specific Regulatory Programs in Real PBMC Data

Applying scRegulate to the PBMC scRNA-seq data revealed distinct TF activity patterns across 8 immune cell types (Figure 4A). In contrast, the latent representation of gene expression based on principal component analysis (PCA) resulted in overlapping clusters, demonstrating that scRegulate’s regulatory-centric TF activity provides a comparable and biologically interpretable representation of cell states (Figure S7). Differential analysis of TF activity identified significantly active TFs specific to the immune cell types (Figure 4B). PAX5 is a well-established master regulator of B cell lineage commitment and maintenance, orchestrating B cell identity through repression of alternative lineage genes (Medvedovic *et al*., 2011). Additionally, BCL6 and IRF4 were recovered, both essential for germinal center formation and plasma cell differentiation, respectively (Ochiai *et al*., 2013). In CD4 T cells, canonical regulators GATA3 and BATF were identified, consistent with their established roles in Th2 differentiation and effector T cell programming (Schraml *et al*., 2009). CD8 T cells exhibited high activity of RUNX3, EOMES, STAT4, STAT5A, and TBX21, which collectively orchestrate cytotoxic T cell lineage specification, expansion, and effector function (Kallies *et al*., 2009). For NK cells, ID2, EOMES, and TBX21critical for cytotoxic development and lineage commitment were among the top-scoring TFs (Gordon *et al*., 2012). In the myeloid lineage, SPI1 (PU.1) was consistently identified across monocytes and dendritic cells, reflecting its central role in myeloid fate specification (Heinz *et al*., 2010). CEBPB and CEBPE further supported monocyte identity through their roles in granulocyte-monocyte differentiation. The recovery of these hallmark TFs provides independent support for the accuracy and interpretability of our inferred TF activity profiles (Figure S8).

**Figure 4.**
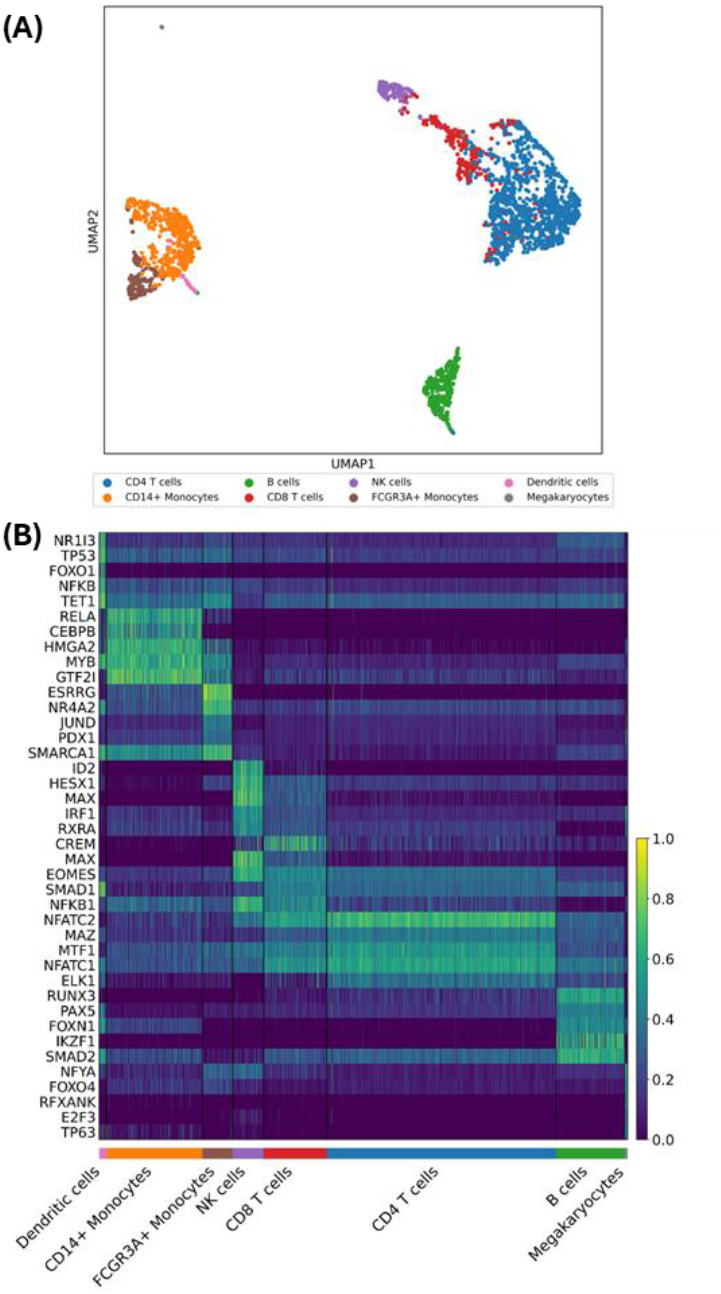
Application of scRegulate in PBMC dataset. **(A)** UMAP visualization of the TF activities demonstrate the cell type heterogeneity, revealing eight distinct cell types. **(B)** Heatmap showcasing the top 5 differentially active TFs per cell type, including TFs well-known for regulating their respective cell types.

Hierarchical clustering of inferred GRNs for the eight cell types further supported the biological relevance of scRegulate’s predictions (Figure S9). For example, CD4 and CD8 T cell subtypes clustered together, reflecting their shared lineage, while monocyte subsets grouped with dendritic cells, mirroring their myeloid origins. Notably, megakaryocytes formed a distinct cluster, highlighting their unique regulatory program separate from immune cells. To explore the structure of these inferred GRN networks, we examined a few DA TFs with their top 3 target gene interactions across cell types (Figure S9A). Key co-regulatory targets emerged, for example, CEBPA, and JUND co-regulate inflammatory genes in monocytes and PAX5 and POU2F2 activating B-cell lineage genes. Functional enrichment analysis confirmed that the inferred top 1% targets for the most differentially active TFs were highly relevant to their associated cell identity (Figure 5). Finally, a TF–TF co-regulatory network revealed that cell-type-specific TFs form tightly connected regulatory modules, with T-cell-associated TFs grouped together, myeloid TFs forming distinct regulatory units, and megakaryocyte TFs positioned independently (Figure S9), as described in Section 2.4.2.

**Figure 5.**
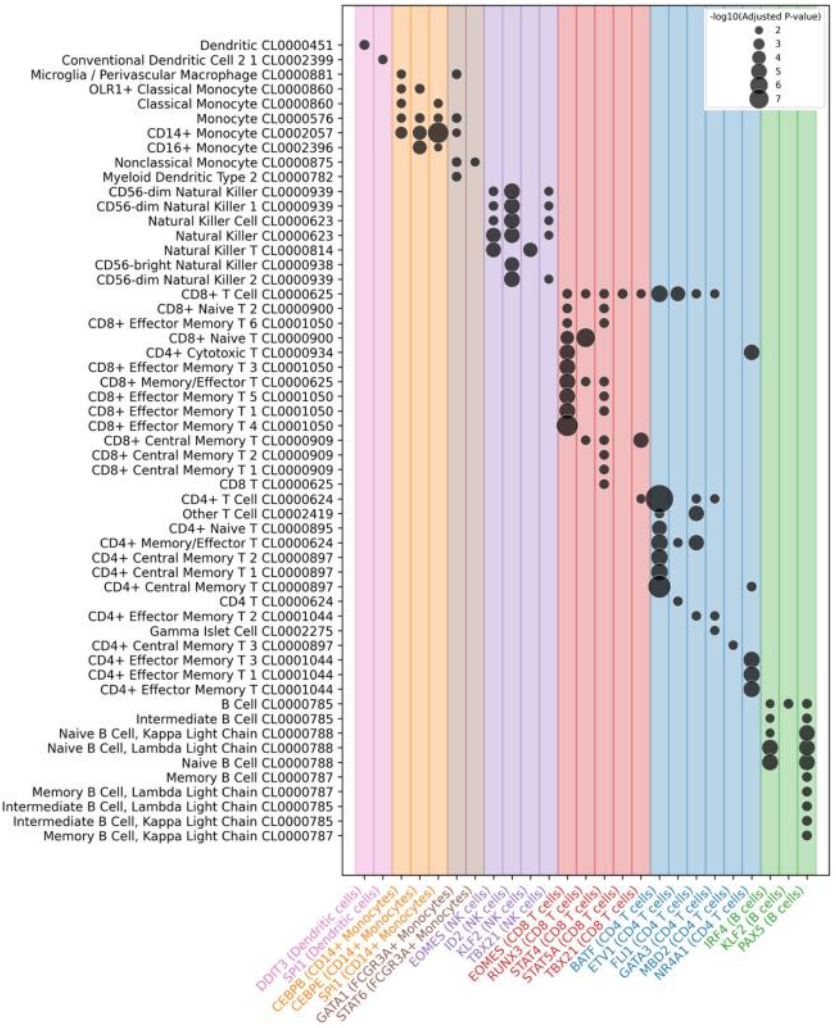
Insights into Cell-Type-Specific Regulatory Networks and Functional Enrichment. Enriched GO terms (Azimuth Human Cell Atlas) based on the top 1% target genes of differentially active TFs from cell type-specific GRN, aligning with cell-type identities.

Overall, scRegulate provides a biologically consistent and interpretable map of TF activity and regulatory interactions in PBMCs. By identifying key regulators, mapping TF-target interactions, and revealing higher-order regulatory relationships, scRegulate provides deeper insights into cell-type-specific transcriptional programs in single-cell data.

## 4 Discussion and conclusions

We developed scRegulate, an interpretable VAE that infers TF activities and GRNs from scRNA-seq data. By embedding TF-gene relationships in the neural network, scRegulate infers cell-type-specific GRNs, predicts novel targets, and estimates per-cell TF activity from the learned representation. Across multiple benchmarks, scRegulate consistently achieve superior or comparable accuracy while being significantly faster than the baseline methods. Beyond inference, scRegulate also supports a wide range of downstream analyses, including clustering cells based on TF activity, identifying differentially active TFs across cell types, and constructing TF-TF coregulatory networks via cosine similarity.

Our benchmarking demonstrated that scRegulate outperformed the existing methods in TF activity-based clustering accuracy across PBMC, lung, heart, and brain datasets compared to decoupleR, SCENIC, BIOTIC, and BITFAM. For the CRISPRi Perturb-seq data, scRegulate accurately predicted negative TF activity changes in knockdown cells, outperforming other methods. These findings highlight scRegulate’s strength in recovering GRNs across diverse contexts. In the PBMC case study, we demonstrated its ability to recover multiple known lineage-defining TFs.

Unlike existing methods, scRegulate jointly models TF activities and GRNs in a cell-type-specific manner. It provides two key outputs: (1) a TF activity representation to identify differentially active TFs and (2) a distinct GRN for each cell population. In contrast, most methods infer TF activity represented by its regulon activity and a global GRN, but do not model cell-type-specific GRNs. In addition, scRegulate leverages all genes that pass basic filtering criteria, overcoming the limitation of approaches that rely on highly variable genes required in other methods. This expansion allows to retain broader regulatory signals for GRN inference. Finally, scRegulate’s ability to infer cell-type-specific GRNs make it unique in offering a high-resolution view of transcriptional regulation at single-cell granularity. Overall, scRegulate fills an important gap between prior-driven inference and deep-learning-based discovery, making it a valuable tool for dissecting single-cell gene regulation.

A major challenge in benchmarking GRN inference is that true regulatory networks are rarely known, especially in higher eukaryotes. Prior studies also highlight the lack of reliable gold standards (Skok Gibbs *et al*., 2022; Kim *et al*., 2024), making synthetic benchmarks essential for ground-truth evaluation. Using the experimental mESC data, we showed that scRegulate achieved better performance compared to one of the best methods (pySCENIC) in much shorter time (Fig. S5). In addition, using the experimental human PBMC data we showed that the predicted TF targets were strongly enriched for cell-type-specific pathways, supporting the biological plausibility of the inferred networks.

A key limitation of scRegulate is its performance dependent on prior GRNs. Our benchmarking (Fig. S10) shows that scRegulate’s performance depends on the networks used for establishing prior knowledge. CollecTRI consistently performs best, likely due to its higher proportion of experimentally validated interactions compared to randomized or uninformative baselines (A–B) and to the broader network (GTRD) or network including text-mined results (TFLink) (C–D). Further hyperparameter tuning may be required in certain datasets where regulatory landscapes differ significantly. As a potential remedy, the prior GRN can be enhanced by incorporating putative TF-gene interactions inferred from motif analysis of chromatin accessibility regions if scATAC-seq data are available under the same biological context (Zandigohar *et al*., 2024). scRegulate’s prior-knowledge guided learning improves interpretability and result stability, making it more robust than purely data-driven methods. Utilization of GPU acceleration overcomes computational bottlenecks for training on large datasets. Expanding scRegulate to model regulatory dynamics across conditions, perturbation responses, would enhance its applicability.

Lastly, we focused on transcription factors as the primary regulators of gene expression because they directly bind regulatory DNA to modulate transcription, supported by extensive prior knowledge such as motif databases and ChIP-seq studies. Other regulators, including chromatin remodelers (Hao *et al*., 2025) and non-coding RNAs (Chen and Kim, 2024), also contribute to transcriptional regulation and can impact GRNs. Our GRN priors do not explicitly model DNA motifs or accessibility, which are key determinants of TF-DNA binding. Future extensions could incorporate DNA-guided features (Xie *et al*., 2025; Seungsoo Kim *et al*., 2024; Gao *et al*., 2022). Studying their roles in GRNs is beyond the scope of the current study, but they could be incorporated into our framework as reliable prior knowledge become available in the future.

## Acknowledgements

We sincerely thank Brandon Lukas for his valuable feedback on this manuscript. This work was supported by the National Institute of Diabetes and Digestive and Kidney Diseases (NIDDK) through the Diabetic Complications Consortium under Grants DK076169 and DK115255. The authors declare no conflicts of interest.

## Data Availability and Supplementary Data

Supplementary data are available at Bioinformatics online. All datasets are publicly available and cited throughout the manuscript. scRegulate is available at https://github.com/YDaiLab/scRegulate, and archived at https://doi.org/10.5281/ze-nodo.17139420, with full reproducibility ensured using a global seed of 42.

## Notes

http://www.github.com/YDaiLab/scRegulate

### Competing Interest Statement

The authors have declared no competing interest.

### Summary of Updates

This revised version incorporates all updates made during peer review and matches the manuscript accepted (in press) at Bioinformatics. We added new benchmarking of scRegulate versus pySCENIC on mouse embryonic stem cell scRNA-seq data, with results reflected in the updated Figure S5 and Table S2, and clarified performance comparisons using experimental human PBMC data. We also expanded the related-work section and updated Table S4 to include Dictys and scMTNI as recent dynamic GRN methods. The distinction between synthetic PBMC benchmarking (GRouNdGAN) and experimental human PBMC data has been clarified, and terminology throughout the manuscript now consistently states that scRegulate infers, rather than reconstructs, GRNs.

