## Supplementary Material for "scRegulate: Single-Cell Regulatory-Embedded Variational Inference of Transcription Factor Activity from Gene Expression"

Contents:

**Note 1**: Univariate linear model

**Note 2**: VAE optimization

**Note 3**: GRouNdGAN synthetic scRNA-seq datasets

**Note 4**: Clustering evaluation metrics

**Note 5**: Criteria for performance evaluation of GRN inference

**Note 6:** TF-TF co-regulation and co-regulatory network analysis

**Table S1**: Symbols used in the manuscripts

**Table S2**: Details of public experimental datasets used in scRegulate evaluation

**Table S3**: GRouNdGAN synthetic datasets used for benchmarking

**Table S4**: Comprehensive overview of representative GRN inference methods from transcriptomics data

**Fig. S1**: Benchmarking of TF activities using four datasets

**Fig. S2**: Benchmarking robustness of clustering performance using TF activity representations in dropout events

**Fig. S3**: Evaluation of scRegulate on MTG data

**Fig. S4**: Receiver Operating Characteristic (ROC) and Precision-Recall (PR) curves used for computing AUROC and AUPRC, respectively

**Fig. S5**: Performance of GRN inference using mouse embryonic stem cells (mESCs) scRNA-seq data

**Fig. S6**: Benchmarking TF activity inference using perturb-seq data

**Fig. S7**: Clustering metrics across resolutions for RNA, TF, latent, and scVI embeddings

**Fig. S8**: UMAPs of the top 5 TFs per cell-type

**Fig. S9**: Cell-type-specific gene regulatory and co-regulatory networks

**Fig. S10**. Benchmarking AUROC and AUPRC performance of scRegulate under different prior constructions.

**Computing environment and LLM use**

**References**

**Supplementary Note 1: Univariate linear model**

To initialize $\hat{\mathbf{e}}_{\mathrm{TF}}$, scRegulate employs a Univariate Linear Model (ULM) inspired by (Badia-i-Mompel *et al.*, 2022a), where TF activities are estimated via:

$$\hat{\mathbf{e}}_{\mathrm{TF}}^{\mathrm{ULM}} = \mathbf{t}\left(\sum_{j=1}^{M} \mathbf{w}_{.,j} \mathrm{x}_j\right)$$

where $\mathbf{w}_{.,j}$ represents known regulatory interactions between all the TFs and gene $j$, $\mathbf{x}_j$ denotes gene expression value for the target gene $j$ and $\mathbf{t}(.)$ computes the vector of t-statistics from the regression model. Briefly, this equation estimates TF activities as a weighted sum of all target gene expression values $\mathbf{x}_j$ in the prior GRN, where $\mathbf{w}_{.,j}$ represents the known regulatory weights for each of the $M$ interactions.

**Supplementary Note 2: VAE optimization**

The VAE is trained to minimize the evidence lower bound (ELBO):

$$\mathcal{L}_{ELBO} = \mathbb{E}_{q_{\boldsymbol{\phi}}(\mathbf{z}|\mathbf{x})}[\log p_{\boldsymbol{\theta}}(\mathbf{e}_{\mathrm{TF}}|\mathbf{z})] + \mathbb{E}_{p_{\boldsymbol{\theta}}(\mathbf{e}_{\mathrm{TF}}|\mathbf{z})}[log p_{\boldsymbol{\theta}}(\mathbf{x}|\mathbf{e}_{\mathrm{TF}})] - \beta D_{KL}(q_{\boldsymbol{\phi}}(\mathbf{z}|\mathbf{x}) \parallel p(\mathbf{z}))$$

$p_{\boldsymbol{\theta}}(\mathbf{e}_{\mathrm{TF}}|\mathbf{z})$ maps $\mathbf{z}$ to TF activities, $q_{\boldsymbol{\phi}}(\mathbf{z}|\mathbf{x})$ is the encoder distribution, $p_{\boldsymbol{\theta}}(\mathbf{x}|\mathbf{e}_{\mathrm{TF}})$ is the likelihood function, and $\beta$ controls the strength of the Kullback-Leibler divergence between the encoder distribution and the prior $p(\mathbf{z})$ for latent space regularization.

The total loss is the sum of the Evidence Lower Bound (ELBO) and the GRN regularization loss:

$$\mathcal{L}_{\mathrm{Total}} = \mathcal{L}_{\mathrm{ELBO}} + \mathcal{L}_{\mathrm{GRN}}$$

Here $\mathcal{L}_{\mathrm{GRN}} = \gamma \sum_j \left\|\mathbf{w}_{.,j}\right\|_1$, where $\gamma$ is the regularization parameter linearly scheduled over training epochs, and the sum is taken over all elements in $\mathbf{w}_{.,j}$, the vector of regulatory weights for gene $j$. The optimizer used for training is Adam, with an adaptive learning rate scheduler that dynamically adjusts based on validation loss. The initial learning rate is set to $\eta_0$ and is reduced according to:

$$\eta^{(n+1)} = \eta_0 \cdot \frac{1}{1 + \lambda t}$$

where $\lambda$ controls the rate of decay, and $t$ represents the iteration. Gradient clipping is applied with a maximum norm of 0.5 to stabilize training.

Unlike standard machine learning approaches that require explicit train-validation-test splits, scRegulate follows common scRNA-seq analysis practices, where benchmarking is performed using external reference datasets rather than separate test splits of the same dataset. To ensure model generalizability and prevent overfitting to noise, we apply an 85%-15% split between the training and validation subsets during model training. The validation loss is monitored during training, and an adaptive scheduler is used to adjust learning rates dynamically based on validation performance. Early stopping is triggered when no significant improvements are observed in validation loss over a set number of epochs, preventing unnecessary training cycles. No explicit test set is used from the same dataset, as benchmarking is performed using independent datasets for GRN inference and TF activity evaluation.

scRegulate supports two modes of fine-tuning: (i) a global unsupervised fine-tuning across all cells, which we used for benchmarking to ensure fairness and avoid label leakage, and (ii) a cell-type–specific mode requiring labels. The latter was applied only in the PBMC case study to extract the most biologically interpretable GRNs aligned with ground-truth cell types and to enable downstream functional enrichment analyses.

A global seed of 42 is used to ensure reproducibility across all computational steps.

**Supplementary Note 3: GRouNdGAN synthetic scRNA-seq datasets**

GRouNdGAN is a reference-based causal implicit generative model that simulates single-cell expression while imposing a user-provided bipartite TF-gene regulatory network (Zinati *et al.*, 2024). The model is trained on an experimental scRNA-seq reference and uses a causal controller to generate TF expression, target generators to produce gene expression conditioned on parent TFs in the GRN, and adversarial learning to match the reference distribution.

**Preprocessing and GRN construction used by GRouNdGAN**

For each reference dataset, cells with nonzero counts in fewer than 10 genes were removed and genes expressed in fewer than 3 cells were discarded. The top 1,000 highly variable genes were selected using the dispersion-based method (Satija *et al.*, 2015), after which counts were library-size normalized to 20,000 per cell. Transcription factors among the highly variable genes (HVGs) were identified using AnimalTFDB 3.0 (Hu *et al.*, 2019), and a TF-gene GRN was inferred with GRNBoost2 (Moerman *et al.*, 2019); in the simulations each gene is regulated by 15 TFs. These steps are reported consistently across datasets in the GRouNdGAN study.

**Synthetic datasets used in our benchmarking**

*PBMC synthetic dataset*. Trained on the 10x Genomics Fresh 68k PBMCs Donor A reference comprising 68,579 PBMCs spanning eleven cell types.

*Tumor-All synthetic dataset*. Trained on a batch-corrected scRNA-seq reference of 136,147 cells from malignant and tumor-microenvironment compartments across 20 follicular lymphoma core-needle biopsies.

*Dahlin synthetic dataset*. Trained on mouse bone marrow HSPCs profiled by 10x Genomics, 44,802 cells (GEO GSE107727), capturing hematopoietic differentiation trajectories.

For PBMC-All, Tumor-All, and Dahlin, we used the publicly released simulated matrices together with the imposed GRNs provided by the authors of GRouNdGAN (include the website that you downloaded from) and analyzed them with our standard preprocessing and evaluation pipeline. See Supplementary Table S3 for details of the datasets used.

**Supplementary Note 4: Clustering evaluation metrics**

The Adjusted Rand Index (ARI) measures clustering agreement while accounting for random chance:

$$ARI = \frac{\sum_{ij}\binom{n_{ij}}{2} - \left[\sum_i \binom{n_{i\cdot}}{2} \sum_j \binom{n_{\cdot j}}{2}\right] / \binom{N}{2}}{0.5\left[\sum_i \binom{n_{i\cdot}}{2} + \sum_j \binom{n_{\cdot j}}{2}\right] - \left[\sum_i \binom{n_{i\cdot}}{2} \sum_j \binom{n_{\cdot j}}{2}\right] / \binom{N}{2}}$$

where $n_{ij}$ represents the number of samples assigned to both cluster $i$ and true label $j$, $n_{i.}$ and $n_{.j}$ are the row and column sums of the contingency table, and $N$ is the total number of samples.

The Normalized Mutual Information (NMI) evaluates clustering similarity based on information theory:

$$NMI = \frac{2I(U,V)}{H(U) + H(V)}$$

where $I(U,V)$ is the mutual information between predicted clusters $U$ and true labels $V$, and $H(U)$, $H(V)$ are their respective entropies.

The F1-score quantifies the balance between precision and recall in cluster assignments:

$$F1 = \frac{2 \cdot Precision \cdot Recall}{Precision + Recall}$$

where precision and recall are defined as:

$$Precision = \frac{TP}{TP + FP}, Recall = \frac{TP}{TP + FN}$$

For multi-class clustering, the macro F1-score is computed as the average of the F1-scores over all classes:

$$Macro\ F1 = \frac{1}{K}\sum_{k=1}^{K} F1_k$$

where $K$ is the number of cell types (clusters).

**Supplementary Note 5: Criteria for performance evaluation of GRN inference**

GRN inference performance is evaluated using AUROC and AUPRC, computed via numerical integration:

$$AUROC = \int_0^1 TPR \ \ dFPR \approx \sum_{i=1}^{1000} (FPR_i - FPR_{i-1})TPR_i$$

Here, TPR is the true positive rate, FPR is the false positive rate, and $N$ (e.g., 1000) is the number of discrete steps used in the numerical approximation.

$$AUPR = \int_0^1 Precision \ \ dTPR \approx \sum_{i=1}^{1000} (TPR_i - TPR_{i-1})Precision_i$$

This computes the area under the precision-recall curve, where the integration is approximated numerically over $N$ steps.

To compare the similarity of inferred GRNs across cell types, Pearson correlation is computed between TF-target interaction matrices:

$$\mathrm{r}_{\mathbf{W}^{(c_1)},\mathbf{W}^{(c_2)}} = \frac{\sum_{i,j}\left(\mathrm{w}_{i,j}^{(c_1)} - \bar{\mathbf{W}}^{(c_1)}\right)\left(\mathrm{w}_{i,j}^{(c_2)} - \bar{\mathbf{W}}^{(c_2)}\right)}{\sqrt{\sum_{i,j}\left(\mathrm{w}_{i,j}^{(c_1)} - \bar{\mathbf{W}}^{(c_1)}\right)^2}\sqrt{\sum_{i,j}\left(\mathrm{w}_{i,j}^{(c_2)} - \bar{\mathbf{W}}^{(c_2)}\right)^2}}$$

where $\mathbf{W}^{(c_1)}$ and $\mathbf{W}^{(c_2)}$ are the inferred GRN weight matrices for two different cell types, and $\overline{\mathbf{W}}^{(c)}$ represents the mean regulatory weight (across all cells and genes of cluster $c$) per GRN.

**Supplementary Note 6: TF-TF co-regulation and co-regulatory network analysis**

TF-TF co-regulation similarity is assessed by computing cosine similarity across rows of $\mathbf{W}$, both within and across cell types, focusing on a biologically relevant subset of TFs. We first selected a few top differentially active TFs from each cell type along with their corresponding GRN weight vectors. We then computed the cosine similarity between each pair of these TFs, both within the same cell type and across different cell types, to quantify the degree of co-regulation. More specifically, assume we have two GRN vectors $\mathbf{w_1}$ and $\mathbf{w_2}$, representing the regulatory weight profiles of a TF from two different cell types (or conditions); The cosine similarity between these two vectors is given by:

$$\text{Cosine Similarity}(\mathbf{w_1}, \mathbf{w_2}) = \frac{\mathbf{w_1} \cdot \mathbf{w_2}}{\|\mathbf{w_1}\|_2 \ \|\mathbf{w_2}\|_2}$$

where $\mathbf{w_1} \cdot \mathbf{w_2}$ is the dot product of the two vectors and $\|\mathbf{w}\|_2$ is the Euclidian norm of a vector $\mathbf{w}$. This analysis allowed us to identify TFs that share similar regulatory profiles and to reveal potential shared regulatory mechanisms across cell types.

Enrichment was evaluated using the Azimuth Cell Types 2021 gene sets (Stuart *et al.*, 2019), applying an adjusted p-value threshold of 0.05 and selecting the top 1% of targets. Relevant GO terms were curated for downstream analyses.

**Table S1**: **Symbols used in the manuscript**

| SYMBOL | MEANING / DESCRIPTION |
|---|---|
| $\mathbf{x}$ | Input gene expression vector for a cell |
| $\mathbf{z}$ | Latent representation (cell embedding in VAE) |
| $\boldsymbol{\mu}(\mathbf{x}), \boldsymbol{\sigma}(\mathbf{x})$ | Mean and standard deviation output of encoder |
| $\boldsymbol{\varepsilon} \sim \mathbf{N}(\mathbf{0}, \mathbf{1})$ | Random noise for reparameterization trick |
| $\hat{\mathbf{e}}_{\mathbf{TF}}$ | Estimated TF activity representation |
| $\mathbf{e}_{\mathbf{TF}}^{\mathbf{ULM}}$ | ULM-based TF activity initialization |
| $\mathbf{p}_{\boldsymbol{\theta}}(\mathbf{e}_{\mathbf{TF}}\|\mathbf{z})$ | Data-driven estimate of TF activity from decoder |
| $\boldsymbol{\alpha}$ | Scheduling parameter balancing prior-driven vs data-driven TF activity |
| $\mathbf{W}$ | TF-gene weight matrix (GRN) |
| $\mathbf{w}_{\mathbf{i,j}}$ | The inferred regulatory strength of TF i on target gene j |
| $\mathbf{w}_{\mathbf{i,\cdot}}$ | The vector of regulatory weights (all target genes) for TF i |
| $\mathbf{w}_{\mathbf{\cdot,j}}$ | The vector of regulatory weights (all TFs) for gene j |
| $\hat{\mathbf{x}}$ | Reconstructed gene expression vector |
| $\boldsymbol{\lambda}$ | L1 regularization parameter on GRN weights |
| $\boldsymbol{m_t}$ | Mask factor applied during training epochs |
| **ELBO** | Evidence Lower Bound (loss function term) |
| **AUROC** | Area under ROC curve (GRN evaluation metric) |
| **AUPRC** | Area under precision–recall curve (GRN evaluation metric) |
| **ARI** | Adjusted Rand Index (clustering metric) |
| **NMI** | Normalized Mutual Information (clustering metric) |
| **F1** | Macro F1-score (clustering metric) |
| **LFC** | Log2 fold-change in TF activity (Perturb-seq validation) |
| **TF** | Transcription Factor |
| **GRN** | Gene Regulatory Network |
| **VAE** | Variational Autoencoder |
| **ULM** | Univariate Linear Model |
| RNA-seq | RNA sequencing |
| scRNA-seq | Single-cell RNA sequencing |
| scATAC-seq | Assay for Transposase-Accessible Chromatin sequencing |
| PBMC | Peripheral Blood Mononuclear Cells |
| GO | Gene Ontology |
| CCC | Cell-Cell Communication |
| TFA | Transcription Factor Activity |

**Table S2**: **Details of public experimental datasets used in scRegulate evaluation**[a]

| Dataset Name | Species | Source Name | Used in | Genes (post-QC) | Cells (post-QC) | GRN Prior Used | # TFs retained | # Targets retained |
|---|---|---|---|---|---|---|---|---|
| Brain | Mus Musculus | Tabula Muris | Figure 2 | 18,631 | 3,401 | collectri_mouse | 264 | 4,768 |
| Heart | Mus Musculus | Tabula Muris | Figure 2 | 13,646 | 624 | collectri_mouse | 222 | 3,748 |
| Lung | Mus Musculus | Tabula Muris | Figure 2 | 15,960 | 5,449 | collectri_mouse | 245 | 4,281 |
| PBMC | Homo Sapiens | 10X -scanpy | Figures 2, 4, 5 | 11,095 | 2,638 | collectri_human | 298 | 3,278 |
| MTG | Homo Sapiens | Allen Brain | Figure 2 | 36,601 | 137,303 | collectri_human | 459 | 5,981 |
| Dixit[b] | Homo Sapiens | Perturb-seq | Figure 3B | 18,531 | 9,439 | collectri_human | 241 | 4,213 |
| mESC | Mus Musculus | Tran et al.[c] | Figure S5 | 3,324 | 6,621 | collectri_mouse | 119 | 2,274 |

[a]In our analysis, we did not restrict to a fixed number of highly variable genes in each dataset. Instead, we used all genes that passed the thresholding and filtering steps during preprocessing.

[b]This perturb-seq dataset contains 10 TFs, each TF was targeted by multiple single-guide RNAs (gRNAs), and each gRNA was sequenced with both forward and reverse reads (ref). To maximize robustness, we combined all gRNAs corresponding to the same transcription factor across their forward and reverse reads. To represent the perturbation of a given transcription factor, we identified cells that expressed at least two distinct gRNAs targeting the same TF (e.g., two different guides against CREB1). These were labeled as double knockdowns (e.g., CREB1 + CREB1) to enrich for stronger functional knockdown effects and reduce noise from single gRNA inefficiency. We included all available transcription factors with double knockdowns, namely ELK1, IRF1, EGR1, GABPA, E2F4, NR2C2, CREB1, ELF1, and ETS1. Cells with detected expression of only intergenic control gRNAs (non-targeting controls) were treated as baseline reference populations.

[c] The 3 gold standard GRNs corresponding to the scRNA-seq dataset of mESCs (Tran *et al.*, 2019) used for benchmarking are curated by McCalla *et al.*, 2023.

**Table S3**: **GRouNdGAN synthetic datasets used for benchmarking*.**

| Dataset Name | Species | Used in | Genes (post-QC) | Cells (post-QC) | GRN Prior Used | # TFs retain ed | # Targets retained | # edges retained |
|---|---|---|---|---|---|---|---|---|
| **PBMC** | Homo Sapiens | Figure 3A | 986 | 99,998 | collectri_human | 24 | 251 | 944 |
| **Tumor** | Homo Sapiens | Figure 3A | 836 | 100,000 | collectri_human | 36 | 276 | 1,556 |
| **Dahlin** | Mus Musculus | Figure 3A | 971 | 100,000 | collectri_mouse | 22 | 323 | 1,001 |

*The table lists the reference species, number of genes and cells after QC, and the GRN priors employed for each dataset.

**Table S4: Comprehensive overview of representative GRN inference methods from transcriptomics data**

| Category | Representative Methods | Notes |
|---|---|---|
| **Co-Expression / Regression Frameworks** | **SCENIC/pySCENIC** (Aibar *et al.*, 2017; Van de Sande *et al.*, 2020)<br>GENIE3 (Huynh-Thu *et al.*, 2010)<br>GRNBoost (Aibar *et al.*, 2017)<br>GRNBoost2 (Moerman *et al.*, 2019)<br>SCENIC+ (Bravo González-Blas *et al.*, 2023)<br>PANDO (Fleck *et al.*, 2023)<br>KIMONO (Ogris *et al.*, 2021; Henao *et al.*, 2023)<br>Inferelator 3.0 (Skok Gibbs *et al.*, 2022)<br>iRafNet (Petralia *et al.*, 2015)<br>NetREX-CF (Wang *et al.*, 2022) | Regression and tree ensemble approaches; SCENIC is combined with motif analysis. |
| **Probabilistic / Bayesian / Graphical Models** | **BITFAM** (Gao *et al.*, 2021)<br>Inferelator (Bayesian modes; Skok Gibbs *et al.*, 2022)<br>BGRMI (Iglesias-Martinez *et al.*, 2016)<br>PriorPC (Ghanbari *et al.*, 2015)<br>Symphony (Bachireddy *et al.*, 2021)<br>BDgraph (Mohammadi and Wit, 2019)<br>Graphical Lasso (Friedman *et al.*, 2008)<br>D-SPIN (Jiang *et al.*, 2025) | Rely on probabilistic inference, graphical models, or Bayesian frameworks; BITFAM tailored for TF activity modeling. |
| **Perturbation / Dynamical Modeling Approaches** | **CellOracle*** (Kamimoto *et al.*, 2023)<br>Dictys (Wang *et al.*, 2023)<br>scMTNI (Zhang *et al.*, 2023)<br>SCODE (Matsumoto *et al.*, 2017)<br>Scribe (Qiu *et al.*, 2020)<br>Inferelator-perturb (Bonneau *et al.*, 2006) | Infer dynamic GRNs or simulate perturbations to model transcriptional responses. |
| **Statistical Aggregation Frameworks** | **decoupleR** (ULM, GLM, VIPER, AUCell, etc.; Badia-i-Mompel *et al.*, 2022b) | Provides unified statistical strategies for TF activity inference. |
| **Neural Network–Based / Deep Learning** | **BIOTIC** (Cao *et al.*, 2025)<br>**scRegulate** (this paper)<br>GRN-VAE (Zhu and Slonim, 2023)<br>GENELink (Chen and Liu, 2022)<br>scGLUE (Cao and Gao, 2022)<br>GRGNN (Wang *et al.*, 2020)<br>scPRINT (Kalfon *et al.*, 2025) | Neural network–based models that embed prior knowledge (GRN/LR) into latent representations. |

*CellOracle builds on regression-based GRN inference but is primarily used as a perturbation/dynamical framework.

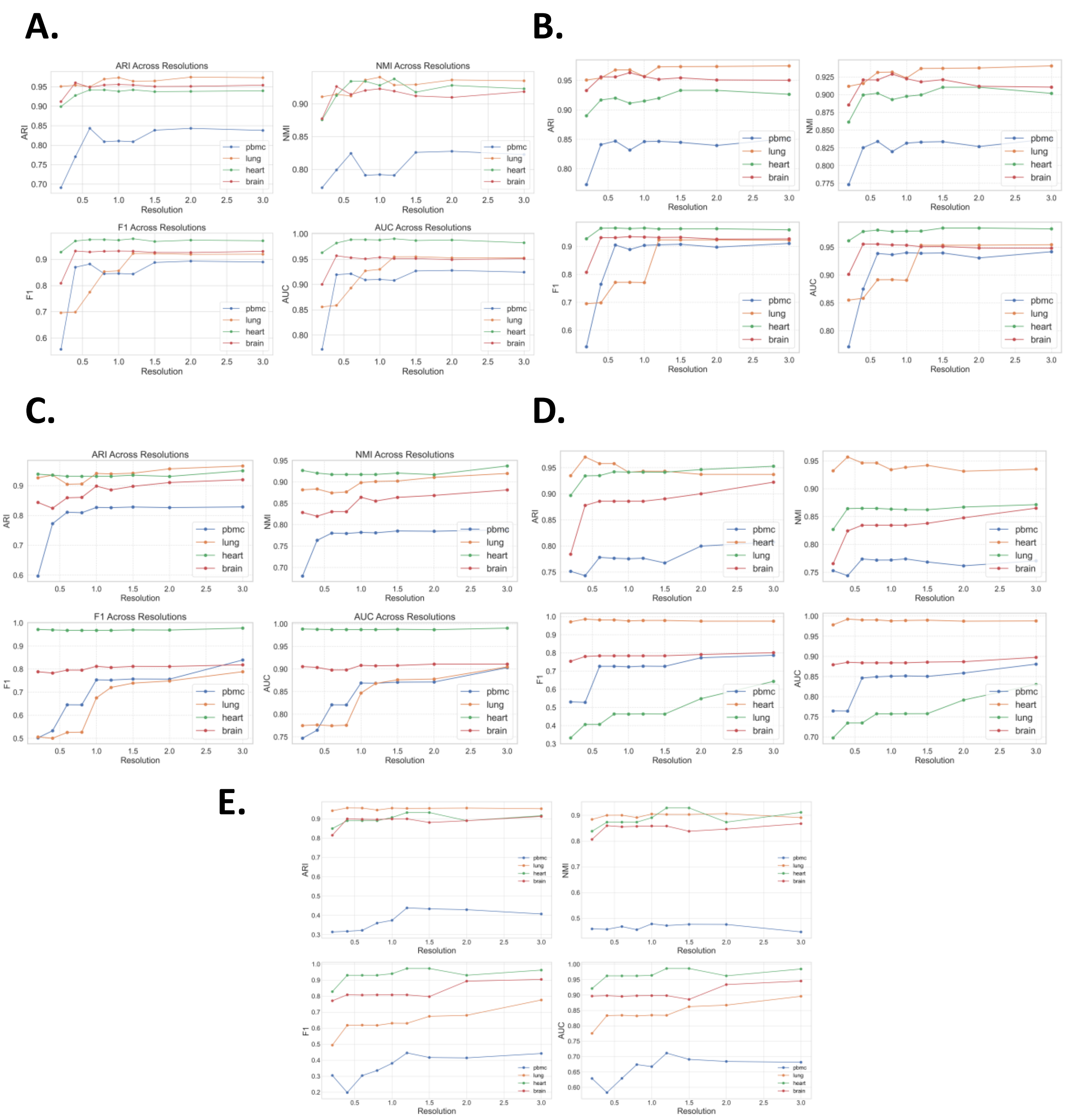


**Fig S1. Benchmarking of TF activities using four datasets.** For a wide range of the Leiden clustering resolutions, we computed different clustering metrics in **(A)** scRegulate, **(B)** decoupleR, **(C)** pySCENIC, **(D)** BITFAM, and **(E)** BIOTIC.

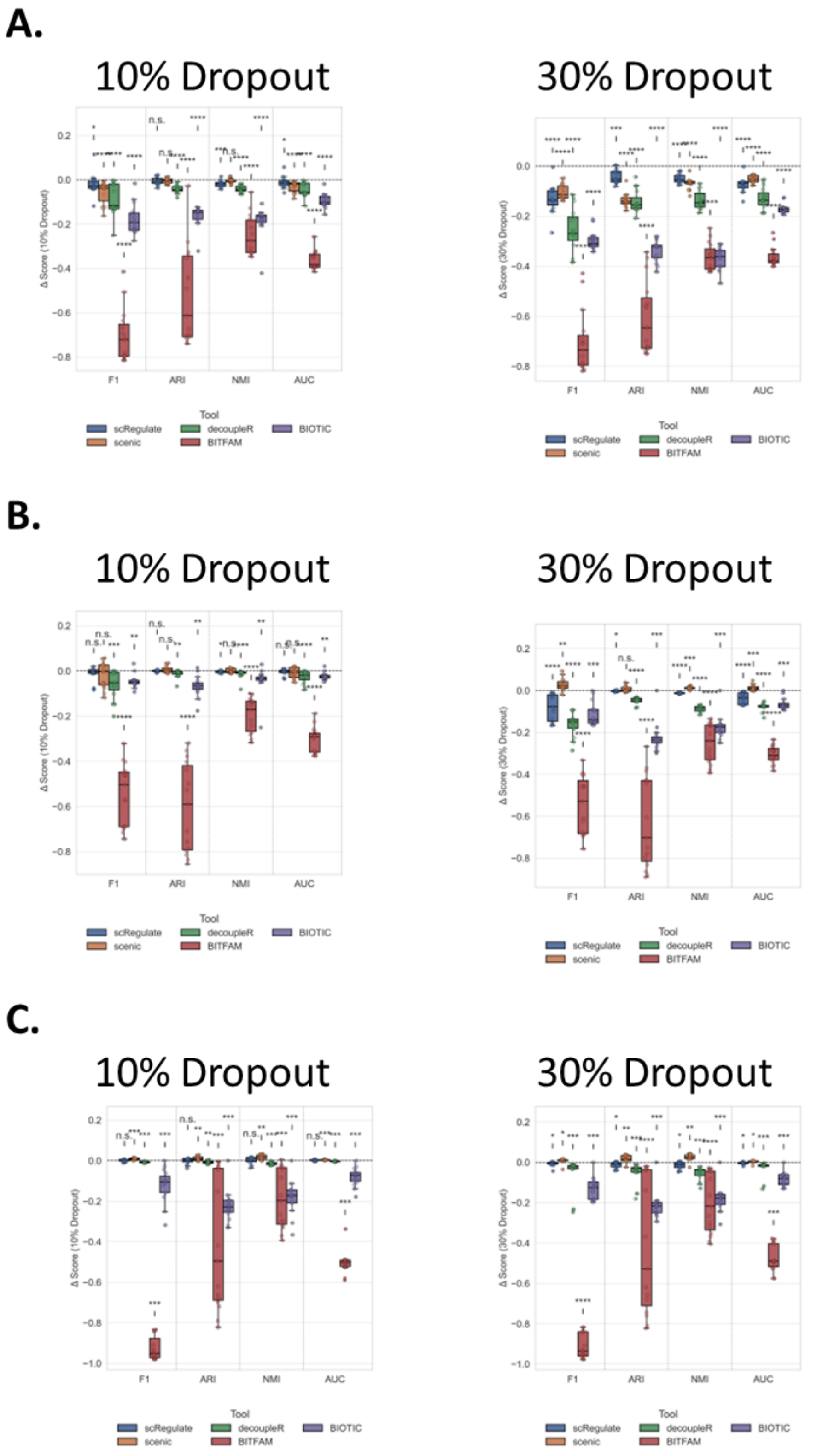


**Fig S2. Benchmarking clustering performance robustness of TF Activity Representations in dropout events.** Each panel shows the drop in different clustering metrics colored by the tool applied for **(A)** PBMC **(B)** Lung, and **(C)** Heart datasets. The drop between the original and noisy datasets in the evaluation metric is indicated by star(s) if statistically significant and otherwise labeled as n.s.

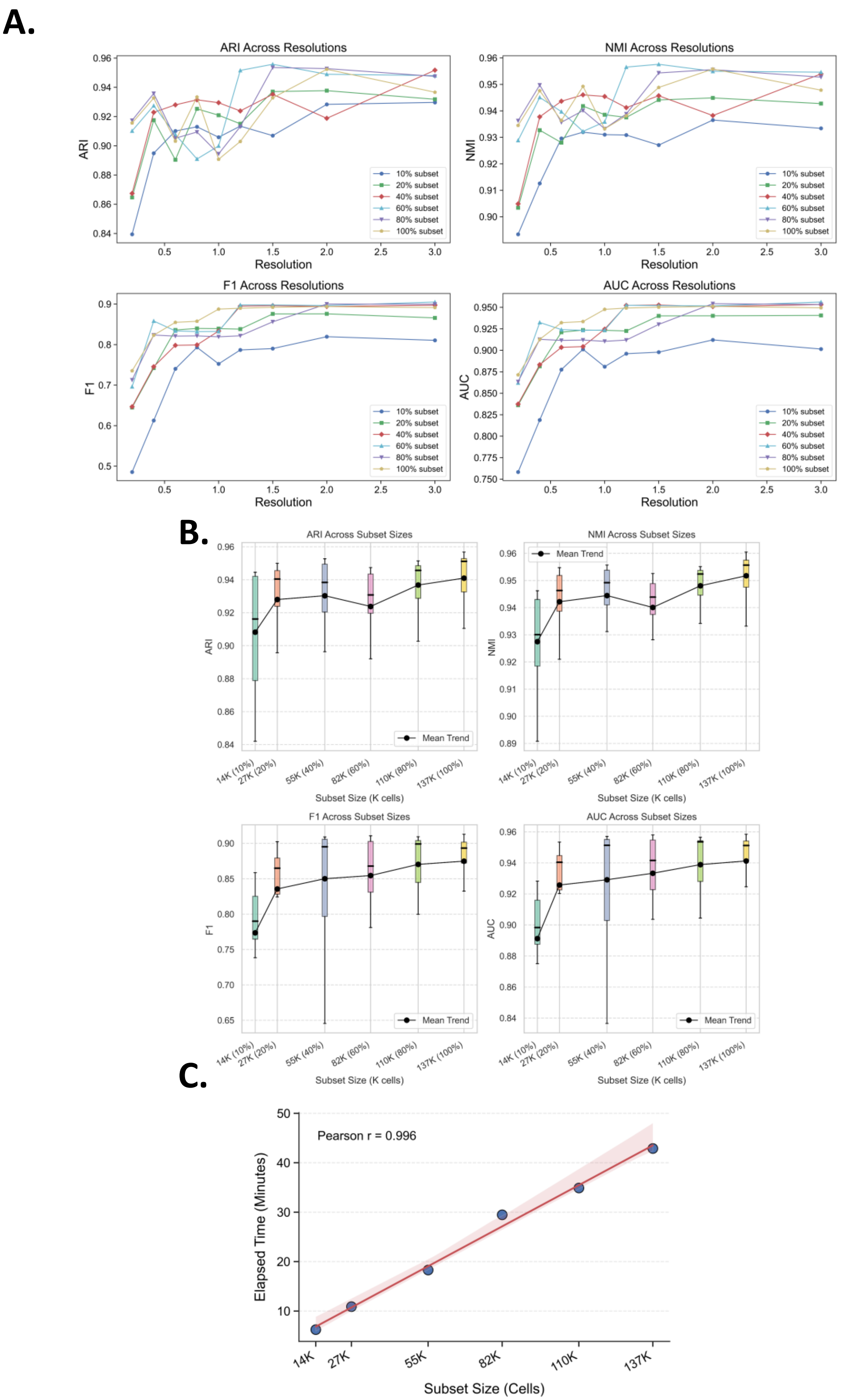


**Fig S3. Evaluation of scRegulate on MTG data. (A)** Shows the clustering evaluation metrics (ARI, NMI, F1 and AUC) for a wide range of the Leiden clusters (resolutions) **(B)** shows the trend of the metrics in (A), and **(C)** Training time vs. dataset size exhibit linear trend $O(n)$.

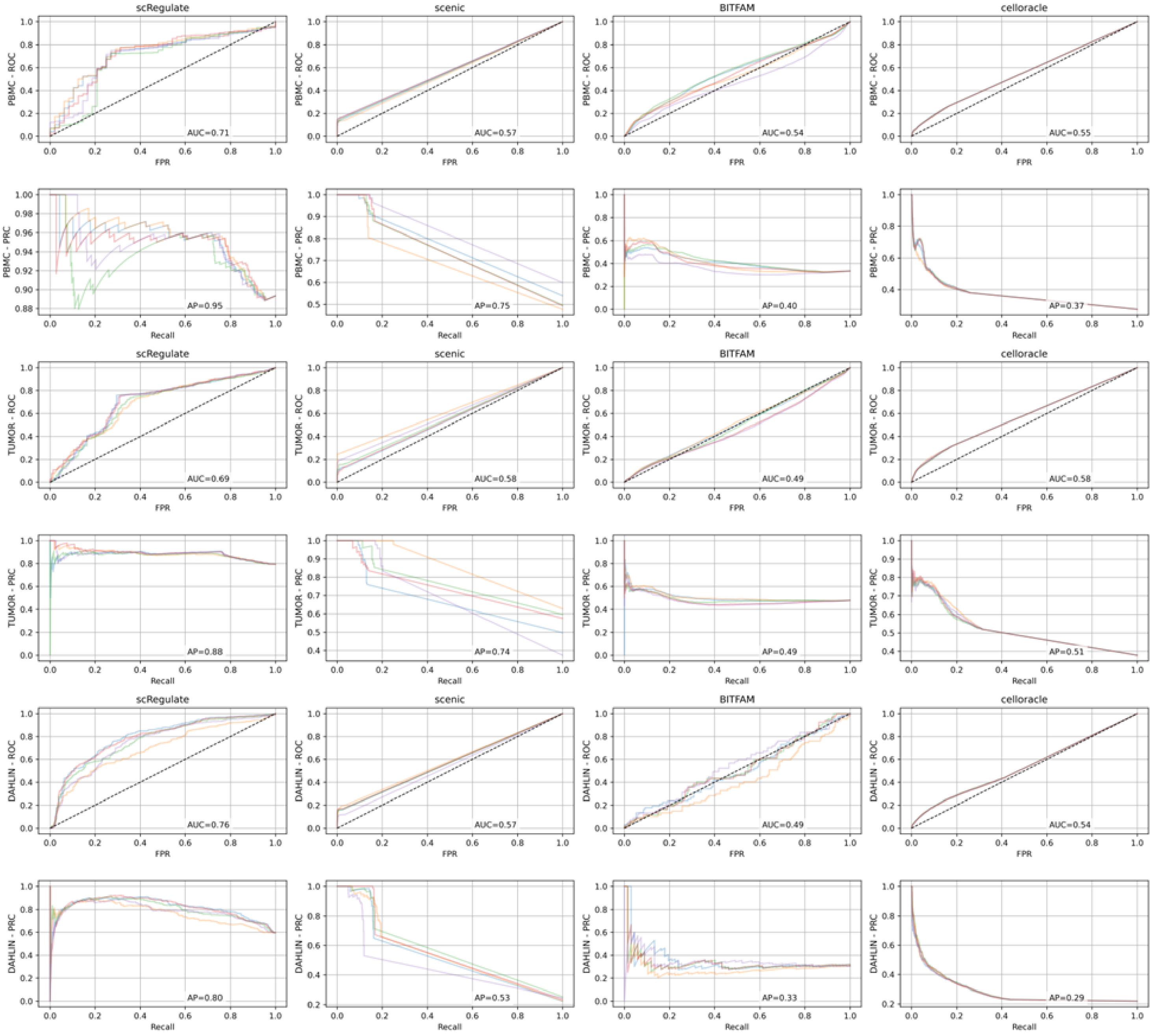


**Fig S4. Receiver Operating Characteristic (ROC) and Precision-Recall (PR) curves used for computing AUROC and AUPRC, respectively. Columns correspond to the (A)** scRegulate, **(B)** pySCENIC, **(C)** BITFAM, and **(D)** CellOracle tools. Rows Are ROC and PRC.

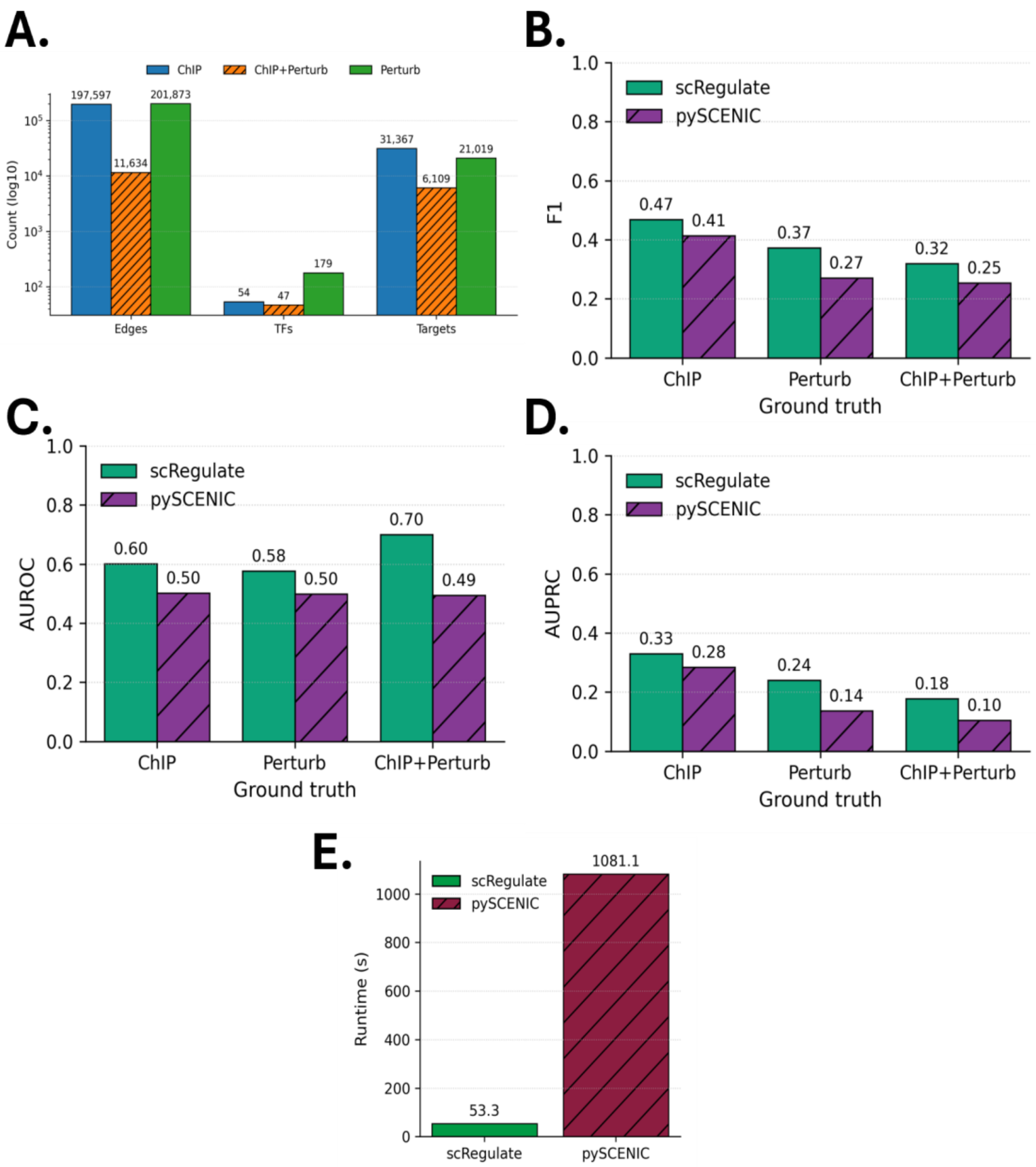


**Fig. S5. Performance of GRN inference using mouse embryonic stem cells (mESCs) scRNA-seq data. (A)** Overlap counts for ChIP, Perturb, and their intersection (ChIP+Perturb) shown for TF-target edges, TFs, and targets in the three gold standard GRNs used in McCalla *et al.*, 2023. ChIP+Perturb TF and target counts reflect unique TFs and genes present in the intersected edge set. Benchmarking metrics are **(B)** F1 at the best threshold, **(C)** AUROC and **(D)** AUPRC values comparing with pySCENIC and scRegulate. **(E)** Run time of scRegulate and pySCENIC on the same computer.

scRNA-seq dataset of mESCs: Tran *et al.*, 2019 (original data includes 3,324 genes × 6,621 cells).

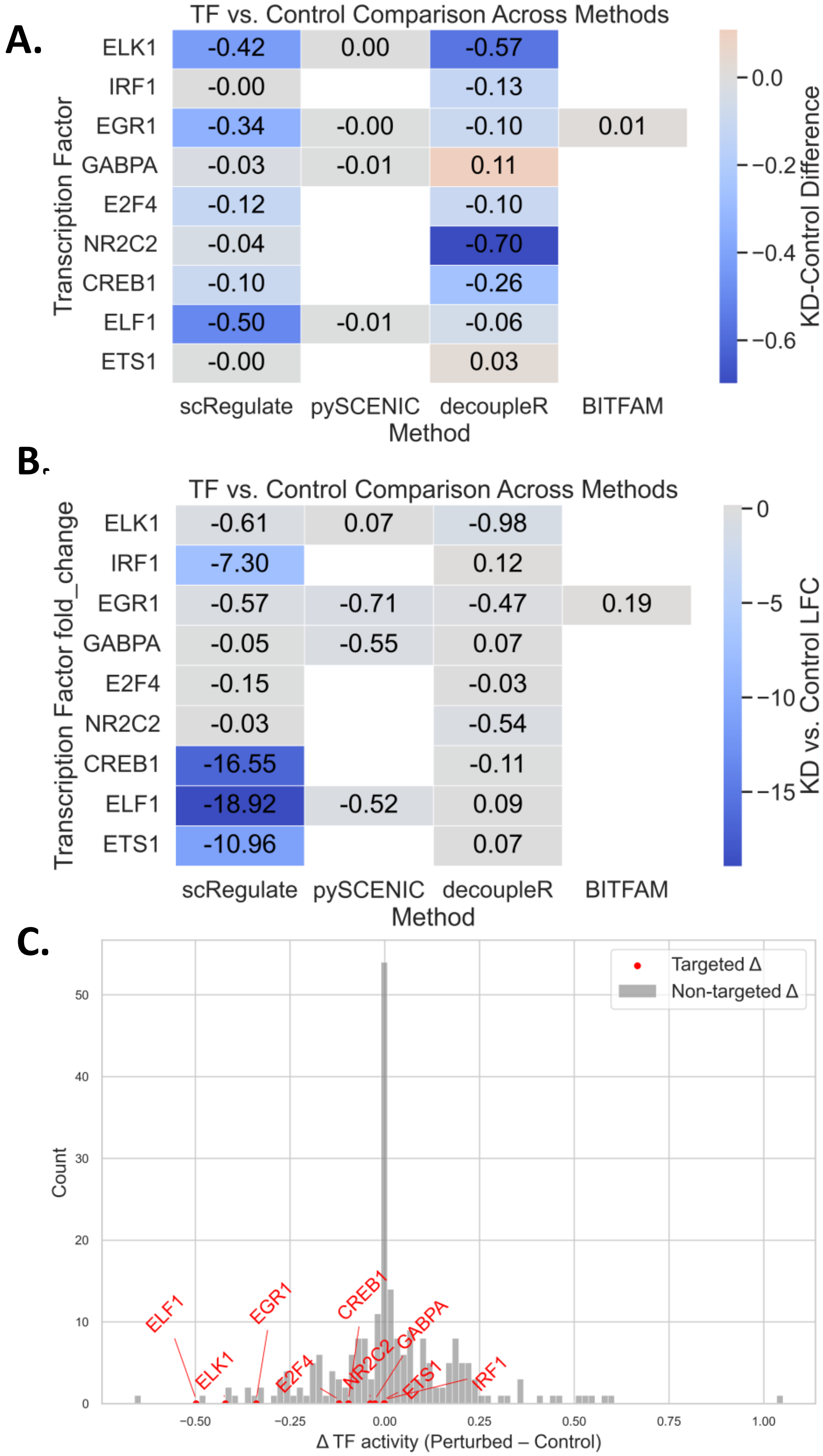


**Fig S6. Benchmarking TF activity inference using Perturb-seq data. (A)** Shows the absolute differences in TF activities between knockout vs. control. **(B)** shows the same comparison but based on log2fold change. **(C)** Distribution of Δ TF activity (perturbed – control) across all TFs. Non-targeted TFs (gray) cluster around zero, while targeted TFs (red) exhibit pronounced negative shifts, indicating that decreases are specific to knockdowns and not due to systematic false positives.

A.

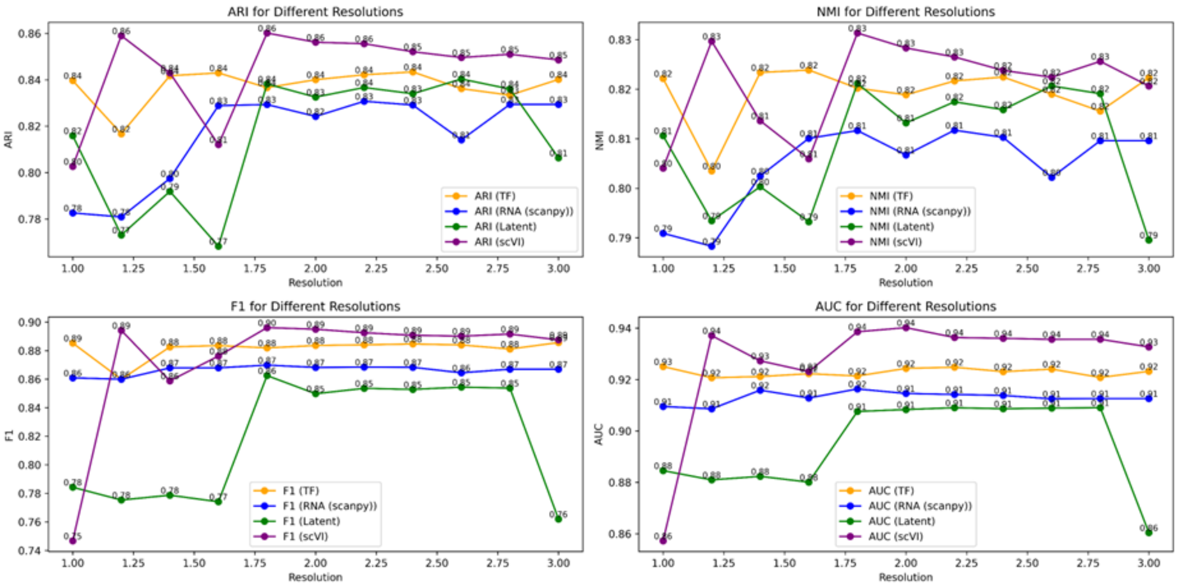


B.

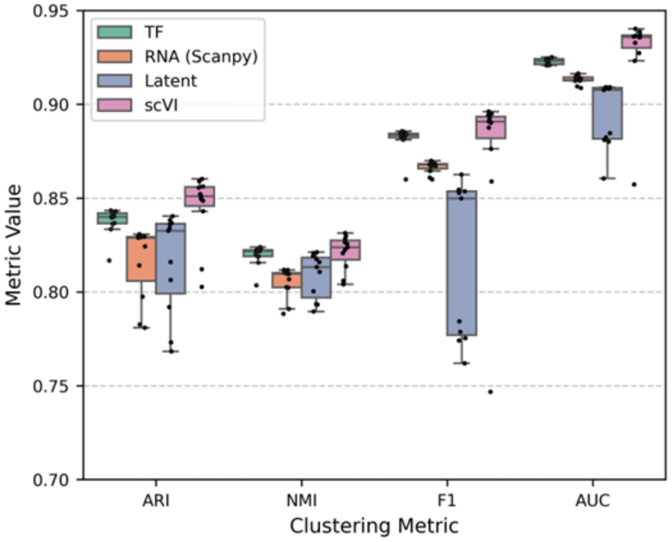


**Fig S7. Clustering metrics across resolutions for RNA, TF, latent, and scVI embeddings. (A)** Post-training evaluation of the TF activity, latent representation, RNA (Scanpy), and scVI. Performance is shown for ARI, NMI, F1, and AUC. TF activities provide a biologically interpretable representation with performance comparable to RNA-based approaches, while scVI serves as a state-of-the-art RNA baseline. Latent embeddings show greater variability, whereas TF activities demonstrate stable and interpretable clustering performance across resolutions. **(B)** scRegulate's nonlinear TF activity representation achieves more stable performance compared to PCA-based RNA reduction, while scVI provides a state-of-the-art RNA baseline.

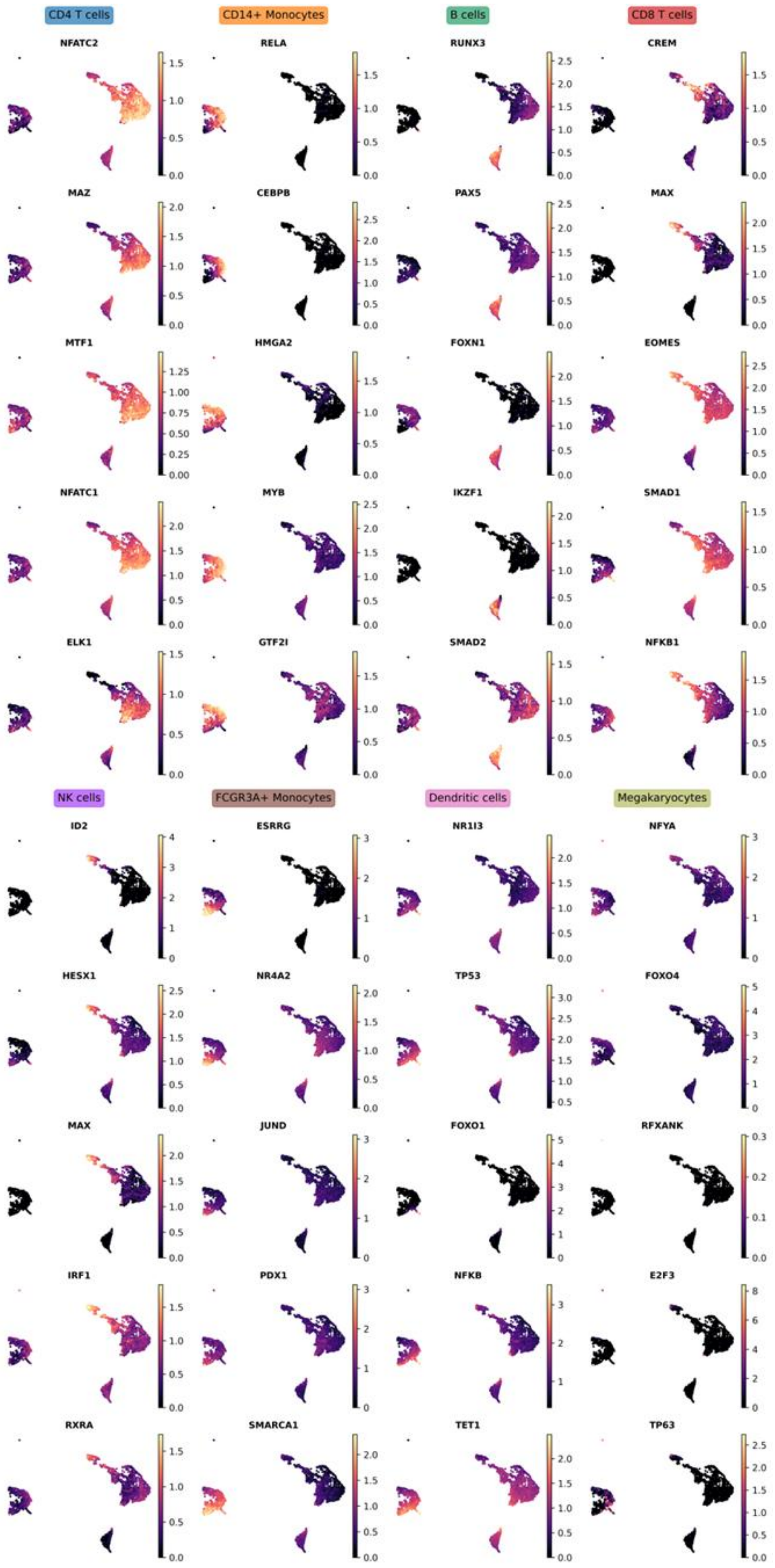


**Fig S8. UMAPs of the top 5 TFs per cell-type** shows scRegulate's capability of inferring cell-type specific TFs.

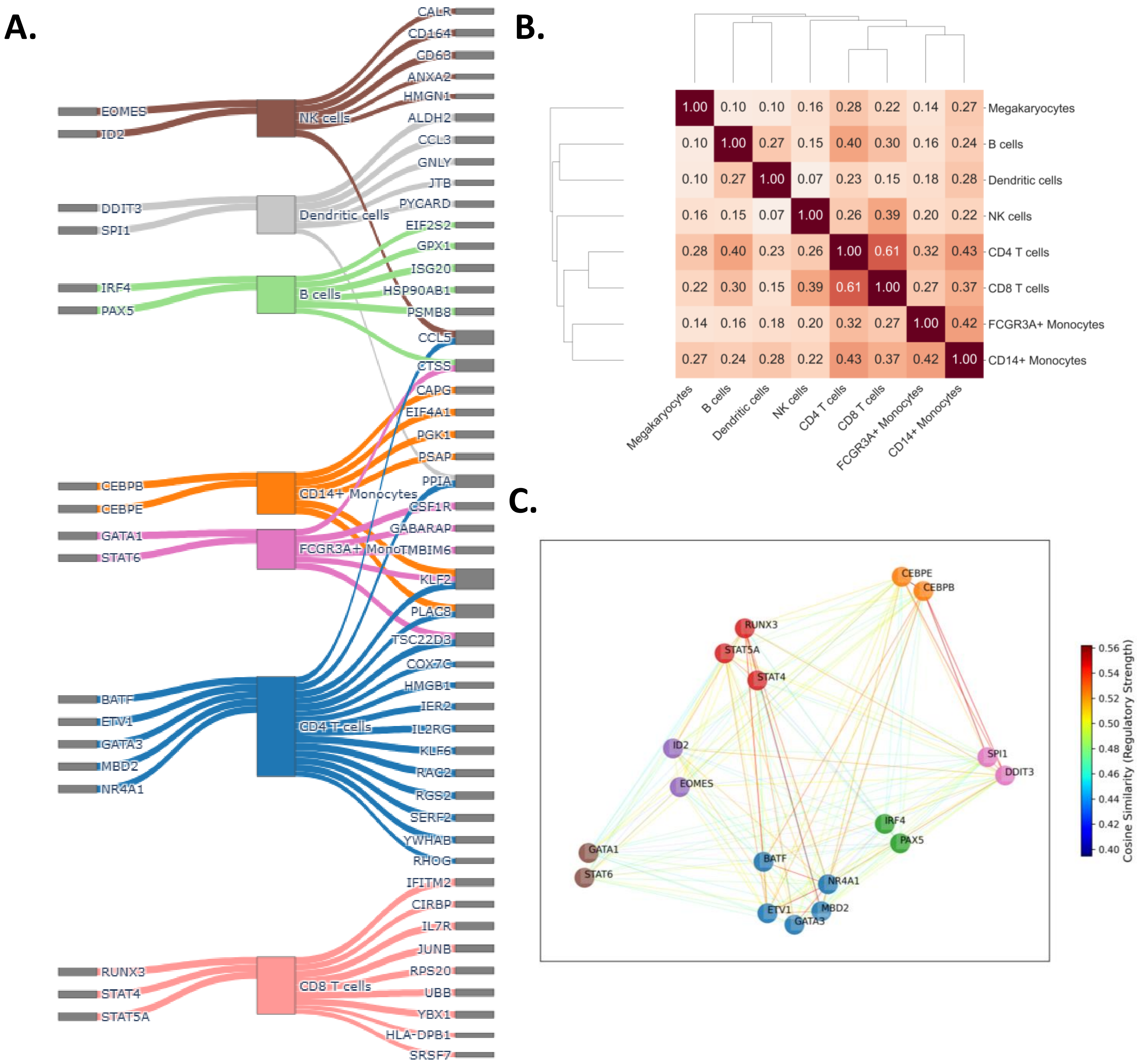


**Fig. S9. Cell-type-specific gene regulatory and co-regulatory Networks. (A)** Sankey plot showing top differentially active transcription factors (TFs) (left) and their top three target genes based on the edge weights, highlighting key TF-target interactions. **(B)** Hierarchical clustering of cosine similarities (raised to the power of 8) between cell type-specific GRN matrices reveals biologically meaningful relationships, with T cells clustering closely together, while megakaryocytes stand out with markedly distinct regulatory patterns. **(C)** TF-TF coregulatory network of the TFs in panel C, with edge colors representing cosine similarity of target gene profiles and node colors representing cell-types. The GRN profiles were extracted from cell-specific networks.

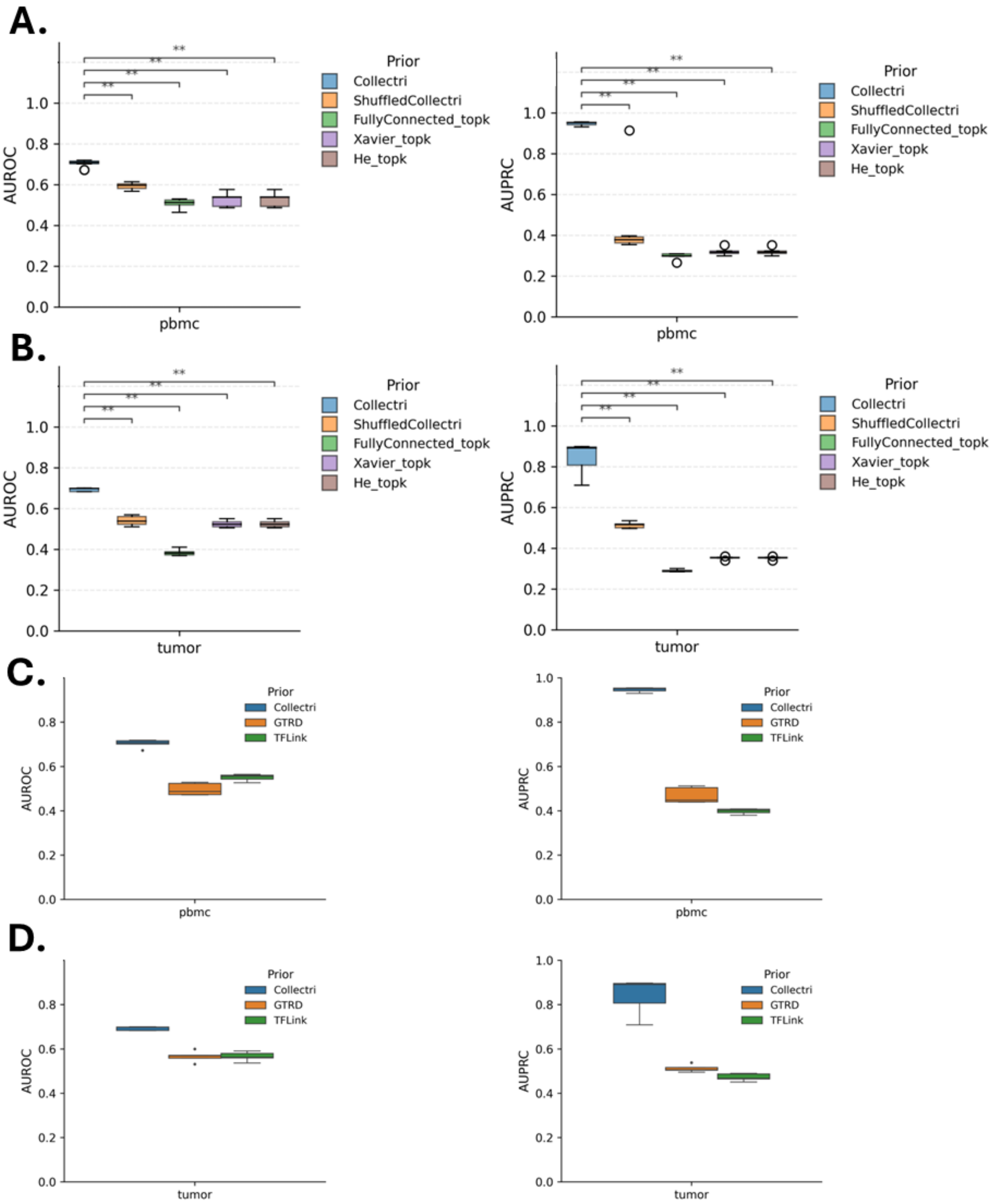


**Fig. S10. Benchmarking AUROC and AUPRC performance of scRegulate under different prior network constructions. (A-B)** Comparison of CollecTRI against randomized (ShuffledCollecTRI), and uninformative (FullyConnected, Xavier, He) using the PBMC **(A)** and tumor **(B)** datasets. CollecTRI consistently outperforms all alternative priors, with statistical tests (Mann-Whitney U, Holm correction) showing significantly higher performance ($p < 0.01$, annotated with **). **(C-D)** Comparison of CollecTRI with alternative public priors (GTRD, TFLink) across PBMC **(C)** and tumor **(D)** datasets further confirms CollecTRI as the strongest baseline across both AUROC and AUPRC metrics.

**Computing environment and LLM use**

We performed all analyses on a Linux workstation (Ubuntu 24.04 LTS, kernel 6.14.0-33) equipped with an Intel Core i7-13700K processor, 62 GiB of memory, and an NVIDIA TITAN Xp GPU, using Python 3.12.2 and PyTorch 2.2.0+cu121 (CUDA 12.1).

ChatGPT (OpenAI) was used solely as a tool to refine and optimize code during the development of this study. It did not contribute intellectually to the study design, interpretation of results, or manuscript writing. All code outputs were manually reviewed, validated, and adapted by the authors. No text, figures, or other content in the manuscript was generated by ChatGPT. This usage complies with the acceptable use policy for large language models as defined by the International Society for Computational Biology (ISCB), and this statement is included in accordance with the journal's guidelines.
